# Highly multiplexed spatially resolved gene expression profiling of mouse organogenesis

**DOI:** 10.1101/2020.11.20.391896

**Authors:** T. Lohoff, S. Ghazanfar, A. Missarova, N. Koulena, N. Pierson, J.A. Griffiths, E.S. Bardot, C.-H.L. Eng, R.C.V. Tyser, R. Argelaguet, C. Guibentif, S. Srinivas, J. Briscoe, B.D. Simons, A.-K. Hadjantonakis, B. Göttgens, W. Reik, J. Nichols, L. Cai, J.C. Marioni

## Abstract

Transcriptional and epigenetic profiling of single-cells has advanced our knowledge of the molecular bases of gastrulation and early organogenesis. However, current approaches rely on dissociating cells from tissues, thereby losing the crucial spatial context that is necessary for understanding cell and tissue interactions during development. Here, we apply an image-based single-cell transcriptomics method, seqFISH, to simultaneously and precisely detect mRNA molecules for 387 selected target genes in 8-12 somite stage mouse embryo tissue sections. By integrating spatial context and highly multiplexed transcriptional measurements with two single-cell transcriptome atlases we accurately characterize cell types across the embryo and demonstrate how spatially-resolved expression of genes not profiled by seqFISH can be imputed. We use this high-resolution spatial map to characterize fundamental steps in the patterning of the midbrain-hindbrain boundary and the developing gut tube. Our spatial atlas uncovers axes of resolution that are not apparent from single-cell RNA sequencing data – for example, in the gut tube we observe early dorsal-ventral separation of esophageal and tracheal progenitor populations. In sum, by computationally integrating high-resolution spatially-resolved gene expression maps with single-cell genomics data, we provide a powerful new approach for studying how and when cell fate decisions are made during early mammalian development.

## Introduction

Lineage priming, cell fate specification and tissue patterning during early mammalian development are complex processes involving signals from surrounding tissues, mechanical constraints, and transcriptional and epigenetic changes, which together prompt the adoption of unique cell fates^1–7^. All of these factors play key roles in gastrulation, the process by which the three germ layers emerge, and the body axis is established. Subsequently, the germ layer progenitors, formed during gastrulation, will give rise to all major organs in a process known as organogenesis.

Recently, single-cell RNA-sequencing (scRNA-seq) and other single-cell genomic approaches have been used to investigate how the molecular landscape of cells within the mouse embryo changes during early development. In particular, these methods have provided insights into how symmetry breaking of the epiblast population leads to commitment to different fates as the embryo passes through gastrulation and on to organogenesis^1–3,6–14^. By computationally ordering cells through their differentiation (“pseudotime”), an understanding of the molecular changes that underpin cell type development has been obtained, providing insight into the underlying regulatory mechanisms, including the role of the epigenome. Recently, technological advances have enabled scRNA-seq to be performed alongside CRISPR/Cas9 scarring, thus simultaneously documenting a cell’s molecular state and lineage. Such approaches have been applied to track zebrafish development^15–17^ and more recently mouse embryogenesis^9,18^. Together, these experimental strategies have enhanced our understanding of developmental lineage relationships and the associated molecular changes.

However, to date, single-cell genomics studies of early mammalian development have focused on profiling dissociated populations of cells, where spatial information is lost. Although regions of the embryo have been micro-dissected and profiled using small cell-number RNA-sequencing protocols, these approaches neither scale to later stages of development, where tens of thousands of cells are present within an embryo, nor do they yet provide single-cell resolution, which may be critical given the role of local environmental cues in conditioning cell fate and patterning at these developmental stages^13,19,20^. By contrast, *in situ* hybridization, single-molecule RNA FISH and other related approaches allow gene expression levels to be measured within a defined spatial context. However, these approaches are typically limited to either quantifying expression patterns in broad domains^21,22^ or to studying a limited number of genes in an experiment, thus precluding generation of comprehensive cell-resolution maps of expression across an entire embryo, which is key for understanding complex processes such as gastrulation and organogenesis. Recent technological advances promise to overcome these limitations: approaches that exploit highly-multiplexed RNA FISH^23–28^, sequencing on intact tissues^29–31^, or that hybridize tissue sections to spatially-barcoded microarrays^32,33^ promise to simultaneously profile the expression of hundreds or thousands of genes within single cells whose spatial location is preserved.

Here, using an existing scRNA-seq atlas covering stages of mouse development from gastrulation to early organogenesis^6^ (‘Gastrulation atlas’), we designed probes against a panel of 387 genes and spatially localized their expression in multiple 8-12 somite stage embryo sections using a version of the seqFISH (sequential fluorescence *in situ* hybridization) method modified to allow highly-effective cell segmentation. Assigning each cell in the seqFISH-profiled embryos a distinct cell type identity revealed different patterns of co-localization of cells within and between cell types. Integrating scRNA-seq and seqFISH data enabled the genome-wide imputation of expression, thus generating a complete quantitative and spatially-resolved map of gene expression at single-cell resolution across the entire embryo. To illustrate the power of this resource, we used these imputed data to perform a virtual dissection of the mid- and hind-brain region of the embryo, uncovering spatially resolved patterns of expression associated with both the dorsal-ventral and rostral-caudal axes. Finally, by integrating a second, independent scRNA-seq dataset that characterized cell types within the developing gut tube^2^, we resolved the position of two clusters of cells that were both previously assigned a lung precursor identity using the scRNA-seq data^2^. Our spatial data revealed that these two clusters were exclusively located on either the dorsal or ventral side of the gut tube, with corresponding transcriptional differences indicating that the dorsal cells give rise to the esophagus, while the ventral cells give rise to the lung and trachea.

## Results

### A single-cell spatial expression profiling of mouse organogenesis

We performed seqFISH^10,11^ on sagittal sections from three mouse embryos at the 8-12 somite stage, corresponding to embryonic day (E)8.5-8.75 (Figure 1A-C). The sections analyzed were chosen to correspond as close as possible to the midline of the embryo, albeit some variation along the left-right axis could be observed due to embryo tilt (Figure 1B). In each section we probed the expression of 351 barcoded genes specifically chosen to distinguish distinct cell types at these developmental stages (Supplementary Figure 1; Supplementary Table 1-2). To do this, we exploited a recently published single-cell molecular map of mouse gastrulation and early organogenesis^6^, and determined computationally a set of lowly-to moderately-expressed genes that were best able to recover the cell type identities (Methods; Supplementary Figure 1). Low-to moderately-expressed genes were selected since low overall expression of the library is needed to reduce the optical density of detected transcripts in a cell so that crowding does not prevent single mRNA spots from being resolved reliably.

**Figure 1:**
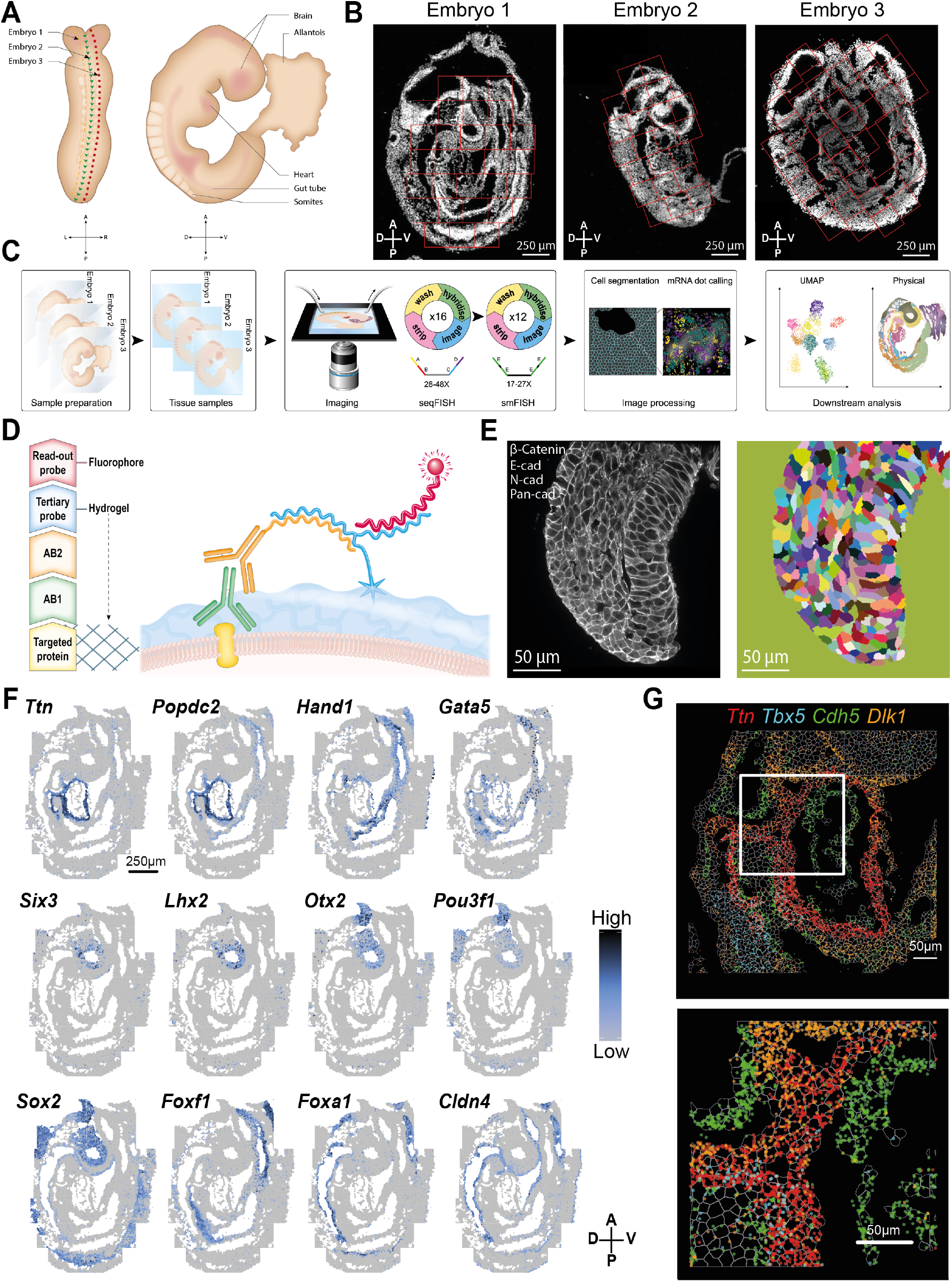
Single-cell spatial transcriptomics map of mouse organogenesis using seqFISH. (A) Illustration of 8-12 somite stage mouse embryo. Horizontal lines indicate estimated position of sagittal tissue section shown in (B). Abbreviations used: D= dorsal, V= ventral, R= right; L= left; A= anterior; P= posterior. (B) Tile-scan of a 20 μm sagittal section of three 8-12 somite stage embryos, stained with nuclear dye DAPI (white). Red boxes indicate selected field of view (FOV), imaged using seqFISH. (C) Illustration of experimental overview for spatial transcriptomics, using seqFISH for 351 selected genes and non-barcoded sequential smFISH for 36 genes. (D) Cell segmentation strategy, using a combination of E-cadherin (E-cad), N-cadherin (N-cad), Pan-cadherin (Pan-cad) and β-catenin antibody (AB; green) staining, detected by an oligo conjugated anti-mouse IgG secondary antibody (orange) that gets recognized by a tertiary probe sequence. The acrydite group (blue star) of the tertiary probe (blue) gets crosslinked into a hydrogel scaffold and stays in place even after protein removal during tissue clearing. The cell segmentation labeling can be read out by a fluorophore-conjugated readout probe (red). (E) Cell segmentation staining of a 10 μm thick transverse section of an E8.5 mouse embryo, using the strategy introduced in (D). Cell segmentation signal was used to generate a cell segmentation mask using Ilastik (right panel). (F) Visualization of normalized log expression counts of 12 selected genes, measured by seqFISH to validate performance. Scale bar 250 μm. (G) Highly resolved ‘digital *in situ’* of the cardiomyocyte marker *Titin* (*Ttn*), *Tbx5*, *Cdh5*, and *Dlk*, colored in red, cyan, green and orange respectively. Dots represent individually detected mRNA spots. Box represents an area that was magnified for better visualization. Scale bars 50 μm.

To obtain a good signal-to-noise ratio for the mRNA spots, we performed tissue clearing to reduce the tissue background signal, as introduced before^26,34^. Briefly, the tissue sections were embedded into a hydrogel scaffold, RNA molecules cross linked into the hydrogel, and lipid and protein removed to achieve optimal tissue transparency for seqFISH (Methods). One consequence of depleting proteins is that delineating the cell membrane, and hence cell segmentation, becomes challenging. To address this, prior to tissue embedding we performed immunodetection for selected surface antigens, Pan-cadherin, N-cadherin, β-Catenin, and E-cadherin, which could in turn be recognized by a secondary antibody conjugated to a unique DNA sequence. We then hybridized a tertiary probe to the DNA sequence of the secondary antibody, which had a unique single-molecular FISH (smFISH) readout sequence and an acrydite group. The acrydite group becomes cross-linked into the hydrogel scaffold and remains in position, even after protein degradation^35^. The unique smFISH readout sequence can subsequently be hybridized with a read-out probe conjugated to a fluorophore, allowing the cell membrane to be visualized (Figure 1D) and enabling segmentation using the interactive learning and cell segmentation tool Ilastik^36^. To validate this strategy, we applied it to a 10 μm thick transverse section of an E8.5 mouse embryo, which confirmed labeling of the cell membrane (Figure 1E; Supplementary Figure 2). Before imaging samples for seqFISH, overall RNA integrity was examined by ensuring co-localization of two *Eef2* probe sets, each detected by a unique read-out probe conjugated to a different fluorophore (Supplementary Figure 2; Supplementary Tables 1 and 3).

Following imaging, the resulting data were segmented as detailed above and individual mRNA molecules were detected by decoding barcodes over the multiple rounds of imaging. To guarantee high sample quality, the first round of hybridization was repeated following all intervening hybridization rounds, allowing for consistency of mRNA signal intensity to be assessed (Supplementary Figure 3). In total, following cell-level quality control, we identified 57,536 cells across three embryos with a combined total of 11,004,298 individual mRNA molecules detected. In the embryo tissue sections, each cell contained on average 196 ± 19.3 (mean ± s.e.) mRNA transcripts from 93.2 ± 6.6 (mean ± s.e.) genes (Supplementary Figure 4), corresponding to an average of 26.6% of all gene’s profiled. The set of genes expressed was not biased towards a specific germ layer, with an average of 21.0% ± 1.1% (mean ± se) genes most associated with a mesoderm identity in the E8.5 Gastrulation atlas being expressed per seqFISH cell, through to 31.6% ± 3.3% (mean ± se) of ectoderm genes.

Next, to confirm the quality of our data, we examined the expression of twelve genes (Figure 1F) with well-characterized expression patterns. As expected, the cardiomyocyte markers *Ttn*^37^ and *Popdc2*^38^ showed the highest expression in the region of the developing heart tube, while *Hand1*^39,40^ and *Gata5*^41^ showed expression in the heart, as well as the more posterior lateral plate mesoderm. Similarly, the expression of four known brain markers, *Six3*^42^, *Lhx2*^43^, *Otx2*^44–46^ and *Pou3f1*^47^ confirmed the strongest expression of these genes in the developing brain. Turning to genes that mark broader territories within the embryo, the neural tube marker *Sox2* showed strong expression in the brain and along the dorsal side of the embryo^48,49^. Additionally, expression of the mesoderm marker *Foxf1* was localized to mesodermal cells outlining the developing gut tube, the lateral plate mesoderm and extraembryonic mesoderm of the allantois^50^. Lastly, two gut endoderm markers *Foxa1*^51^ and *Cldn4*^52,53^ marked the developing gut tube along the anterior-posterior axis of the embryo. The tissue-specific expression profile of these genes was consistent with both the Gastrulation atlas^6^ (Supplementary Figure 4) as well as the broad expression territories defined in the EMAGE database^21^. As a further confirmation of the quality of our data, we confirmed the positional expression profiles of the measured Hox gene family members, which followed the described ‘Hox code’ along the anterior-posterior axis^54,55^ (Supplementary Figure 5). Finally, the high-resolution of seqFISH allows visualization of mRNA molecules at sub-cellular resolution, enabling the generation of high quality digital *in situs* (Figure 1G). Taken together, these analyses demonstrate that we can reliably record the expression profiles of hundreds of genes across an entire embryo cross-section at single-cell resolution.

### Cell type identity and spatial transcriptional heterogeneity

Thus far we have focused on the expression of individual genes. However, the real power of the data derives from the ability to study co-expression of hundreds of genes within their spatial context. To develop this potential, as a first step, we assigned each cell within the seqFISH-profiled embryos a distinct cell type identity using cell type mapping. To make this assignment we integrated each cell’s expression profile from seqFISH with the E8.5 cells from the Gastrulation atlas^6^ using batch-aware dimension reduction and Mutual Nearest Neighbours (MNN) batch correction^56^ (Supplementary Figure 6), before annotating seqFISH cells based on their nearest neighbors in the Gastrulation atlas (Figure 2A; Supplementary Figure 6). We further refined this automated cell type classification by performing joint clustering of both datasets and comparing their relative cell type contribution and gene expression profiles (Supplementary Figure 6; Methods). We observed that the assigned cell type identities were consistent with known anatomy as well as with the expression of distinct marker genes (Figure 1F; Figure 2B-C; Supplementary Figure 7-9).

**Figure 2:**
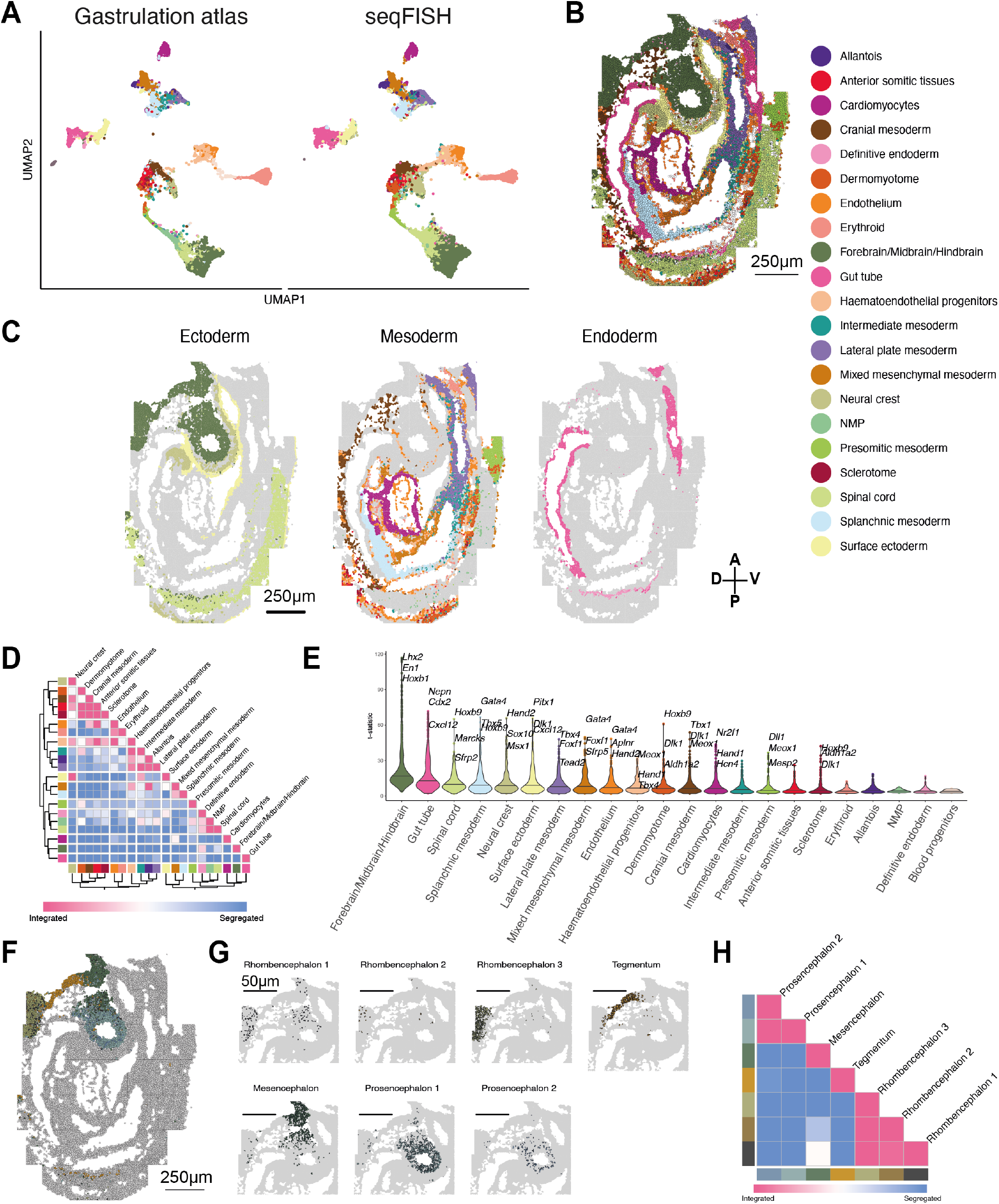
Cell type annotation and neighborhood characterization. (A) Projection of seqFISH spatial and Gastrulation atlas cells in joint reduced dimensional space in order to annotate seqFISH cells based on their nearest neighbors in the mouse Gastrulation atlas. (B) Real position of annotated seqFISH cells in embryo tissue section. Colors represent refined cell type classification. Scale bar 250 μm. (C) Cell type maps separated by the three germ layers (ectoderm, mesoderm, endoderm). Scale bar 250 μm. (D) Cell-cell contact map displaying the relative enrichment towards integration and segregation of pairs of cell types in space. Cell types are clustered by their relative integration with others. (E) Violin plots showing the t-statistic for each gene and cell type corresponding to a measure of the degree of residual transcriptional heterogeneity explained by space. For each cell type selected top genes are labeled. (F) Re-clustering of Forebrain/Midbrain/Hindbrain cell type into 7 spatially distinct clusters. Scale bar 250 μm. (G) Zoom in of the brain region to visualize four major brain regions and seven subclusters identified in (F). Scale bars 50 μm. (H) Cell-cell contact map of brain subclusters in space, ordered roughly anatomically from hindbrain to forebrain.

As an alternative, we performed direct clustering of the seqFISH data, which revealed similar groupings of cells (Supplementary Figure 10), indicating that a small number of carefully-chosen genes can provide enough information to accurately group cells. However, we note that assigning cell type identity using only a small number of marker genes is likely to be less reliable than imputing identity through reference to the Gastrulation atlas.

Next, to study when boundaries between emerging tissue compartments are established in the developing embryo, we statistically quantified whether cells assigned to the same type were spatially coherent within the embryo, as well as determining the extent to which pairs of cell types were co-located (Figure 2D-E, Methods). We used a permutation strategy to evaluate the relative enrichment or depletion of direct cell-cell contact events between each cell type (compared to a random distribution of cell types) resulting in a cell-cell contact map (Figure 2D, Supplementary Figure 11). Certain cell types, such as cardiomyocytes and the gut tube were spatially and morphologically distinct within the embryo, while others, like the endothelium, were interspersed and spread across the entire embryo space.

More generally, while most cell types are characterized using prior knowledge of expression markers and lineage inference, other populations such as the mixed mesenchymal mesoderm represent a cell state rather than a defined cell type. Mesenchyme represents a state in which cells express markers characteristic of migratory cells loosely dispersed within an extra-cellular matrix^57^. This strong overriding transcriptional signature of mesenchyme, irrespective of location, makes it challenging to distinguish which cell types this mixed mesenchymal mesoderm population represents using classical scRNA-seq data. In contrast, our integrated spatial expression map allowed us to resolve five transcriptionally distinct subpopulations (cluster 1-5) that were spatially defined (Supplementary Figure 12; Methods).

Based on its anatomical position overlaying the developing heart, we infer that cluster 1 reflects cells with a cardiac mesoderm and pericardium identity. Clusters 2 and 3 are located in the septum transversum, in the region of the forming hepatic plate and proepicardium. At this developmental stage BMP signaling from the developing heart and FGF signaling from the septum transversum mesenchyme is critical for the induction of hepatic fate specification in the foregut^58,59^. Consistent with this we observed enrichment for BMP signaling in cluster 1 (Supplementary Figure 12). Additionally, in cluster 3 we observed the co-expression of proepicardial markers *Tbx18* and *Wt1*^60,61^ whose deletion results in heart^62^ and liver^63^ defects (Supplementary Figure 12). Our ability to spatially map cluster 3 revealed its position caudal to the forming heart, corresponding with the known location of the proepicardium, thereby allowing us to characterize this cluster. Together, their location and expression profiles indicate that the cells from cluster 2 and 3 will contribute to the hepatic mesenchyme (important for hepatoblast specification) and the proepicardium, respectively. Lastly, cluster 4 and 5 are located toward the body wall, suggesting a somatic mesoderm identity that will contribute to the dermis^64^.

To assess additional, more subtle, spatially-driven transcriptional heterogeneity, we used a linear model to identify genes that show a strong spatial expression pattern within each cell type (Figure 2E; Supplementary Table 4; Methods). This indicated that residual transcriptional heterogeneity in the Forebrain/Midbrain/Hindbrain cluster can be explained by localized patterns of expression, most likely resulting from the presence of regionally-specific developing brain subtypes (Supplementary Table 5). To investigate this further, we performed a focused re-clustering of Forebrain/Midbrain/Hindbrain cells, recovering four major brain subregions and seven subclusters (Figure 2F-G). Cross-referencing spatial location and underlying gene expression signature allowed us to identify subclusters associated with the prosencephalon, mesencephalon, rhombencephalon and the tegmentum (Figure 2G-H; Supplementary Figure 11).

### A single-cell 10,000-plex spatial map of inferred gene expression in the mouse embryo

By design, our seqFISH library allowed us to probe the expression of specific genes associated with cell type identity. Additionally, we directly measured the expression of a number of genes associated with key signaling cascades e.g. Notch^65^ and Wnt^66^. Nevertheless, a full, unbiased, view of the interplay between a cell’s spatial location and its molecular profile, and how this influences development would benefit from measuring expression of the entire transcriptome, something that is not straightforward with existing highly-multiplexed RNA FISH protocols.

To overcome these limitations, we built upon the MNN mapping approach described earlier (Figure 2, Supplementary Figure 6) and inferred the full transcriptome of each seqFISH cell by considering the weighted expression profile of the cells to which it is most transcriptionally similar in the Gastrulation atlas (Figure 3A; Supplementary Figure 13; Methods). To test the integrity of this strategy, for each gene probed in our seqFISH experiment (excluding *Xist*, as it is sex specific), we used the remaining 349 measured genes to map all cells to the Gastrulation atlas and imputed the expression of the withheld gene. To evaluate performance, we calculated, for each gene and across all cells, the Pearson correlation (‘performance score’) between the imputed expression counts and the measured seqFISH expression levels. To estimate an upper bound on the performance score (i.e., the maximum correlation we might expect to observe) we exploited the four independent batches of E8.5 cells that were processed in the scRNA-seq Gastrulation atlas. We treated one of the four batches as the query set and used the leave-one-out approach described above to impute the expression of the 350 genes of interest by mapping cells onto a reference composed of the remaining three batches, before computing the Pearson correlation between the imputed and true expression counts (‘prediction score’; Methods). Computing the ratio of the Performance (seqFISH – scRNA-seq) and Prediction (scRNA-seq – scRNA-seq) scores yields a normalized performance score. Across genes, we observed a median normalized performance score of 0.73 (lower quartile 0.32, upper quartile 1.09) (Supplementary Figure 13), suggesting that our ability to infer gene expression is comparable to what might be expected when combining independent scRNA-seq datasets and providing confidence in our approach.

**Figure 3:**
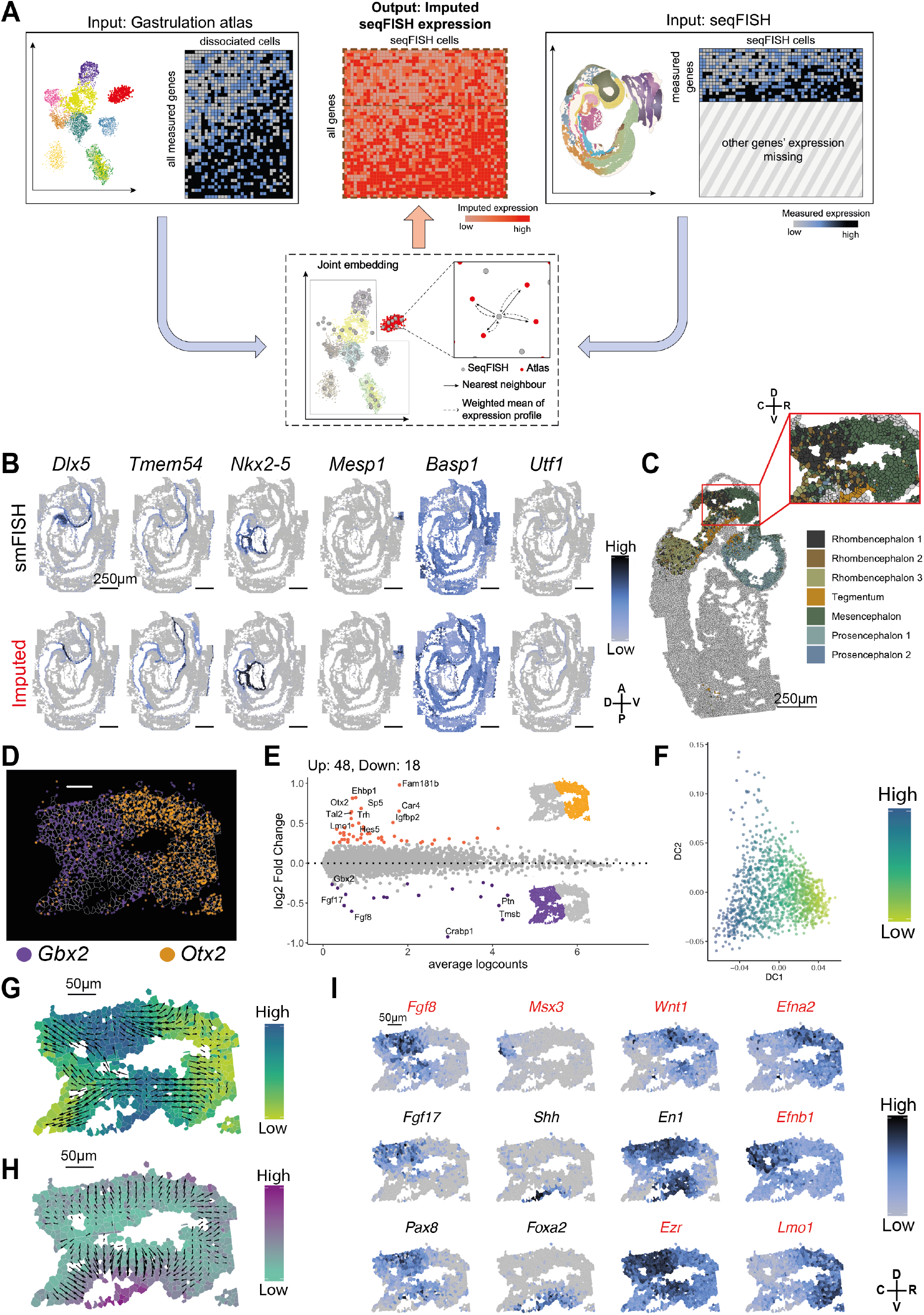
Creating and using a 10,000-plex spatial map. (A) Schematic representation of the imputation strategy. (B) Independent validation of imputation performance by comparing normalized gene expression profiles of selected genes measured by smFISH with the corresponding imputed gene expression profiles. Scale bars 250 μm. (C) Visualization of brain subclusters in embryo 2 and virtual dissection of the midbrain-hindbrain boundary (MHB), highlighted by red rectangle and inset zoom. Scale bar 250 μm. Abbreviations used: C= caudal; R= rostral; D= dorsal; V= ventral. (D) ‘Digital *in situ*’ showing detected mRNA molecules of a mesencephalon and prosencephalon marker *Otx2* (orange dots) and a rhombencephalon marker *Gbx2* (purple dots) to identify the MHB. Scale bar 50 μm. (E) MA (log-ratio and mean average) plot showing differential gene expression analysis between the virtually dissected hindbrain region (orange, 48 genes significantly upregulated, absolute LFC > 0.2, FDR-adjusted P-value < 0.05) and virtually dissected midbrain region (purple, 18 genes significantly upregulated, absolute LFC > 0.2, FDR-adjusted P-value < 0.05) using the imputed transcriptome. (F) Diffusion pseudotime analysis of the virtually dissected region to understand dynamics of gene expression at the MHB. Scatterplot of diffusion-based embedding of virtually dissected cells, displaying diffusion components (DC) 1 and 2. Cell colors correspond to inferred diffusion pseudotime. (G) Spatial graph showing virtually dissected cells colored by inferred diffusion pseudotime, dominated by DC1. Arrow sizes correspond to the magnitude of change of pseudotime value within the region, in the direction from large to small pseudotime values. The highest pseudotime values are observed along the MHB region, smoothly diffusing outward to the midbrain and hindbrain regions. Scale bar 50 μm. (H) Spatial graph showing virtually dissected cells colored by DC2. Arrow sizes correspond to the magnitude of change of DC2 value within the region. The most extreme DC2 values are observed perpendicular to the MHB region, smoothly diffusing outward to the floorplate and roofplate regions. Scale bar 50 μm. (I) Visualization of normalized log expression counts of important regulators of midbrain/hindbrain formation. Gene name in red font indicate imputed expression, while black font indicates measured expression. Scale bar 50 μm.

To further validate our imputation strategy, we used non-barcoded sequential smFISH to measure the expression of 36 additional genes in the embryo sections probed by seqFISH and contrasted the true expression profile with the imputed values (Figure 3B). This independent validation – these smFISH genes were not used in the MNN mapping – confirmed that imputation reliably recovered gene expression profiles (Figure 3B; Supplementary Figure 14-18). For example, we observed a strong overlap between measured and imputed expression for *Dlx5*^67^, an essential and spatially-restricted regulator of craniofacial structures, in the anterior surface ectoderm and first branchial arch. Additionally, we noted that *Tmem54* was inferred to be specifically expressed in the anterior surface ectoderm and along the gut tube, *Nkx2-5*^68,69^ was inferred to be expressed in the developing heart, and *Mesp1* was inferred to be expressed in the posterior presomitic mesoderm (PSM; ^70,71^). Finally, the ubiquitous expression profile of *Basp1* and the absence of expression of the germ line marker *Utf1*^72^ was also recapitulated in the imputed expression maps.

### A whole-genome spatial map allows reconstruction of midbrain-hindbrain boundary formation

To illustrate the utility of the imputed data, we focused on a well-described developmental process that takes place at this embryonic stage – the formation of the midbrain-hindbrain boundary (MHB), also known as the isthmus organizer. The MHB acts as a signaling hub that is essential for patterning of the adjacent midbrain and hindbrain regions by inducing two distinct transcriptional programs via defined signaling cascades (reviewed in ^73–75^). Thus, the MHB presents an important dividing point in the developing brain, functioning both as a signaling center and as a physical barrier of the developing brain ventricles^76^. We observed expression of the mesencephalon and prosencephalon marker *Otx2*^44,77^ and the rhombencephalon marker *Gbx2*^77,78^ in the brain region of all three embryos, albeit the sagittal section for embryo 2 appeared to capture this region most comprehensively (Supplementary Figure 19). Focusing on this region of embryo 2, we used expression of *Gbx2* and *Otx2* to identify the precise boundary between the two subclusters (Figure 3C-D). Subsequently, we virtually dissected the *Otx2* positive midbrain region and the *Gbx2* positive hindbrain region (Supplementary Figure 19) and performed a differential expression analysis (using the imputed expression profiles) to identify additional genes that distinguish the two regions (Figure 3E). This identified 66 genes (FDR-adjusted P-value < 0.05; Absolute log fold change > 0.2) with spatially distinct expression profiles between the two regions (Supplementary Table 6).

To further understand the spatial distribution of gene expression at the MHB, we investigated whether further local differences in spatial expression patterns were present. Using a diffusion-based transcriptional embedding^79^, we observed smoothness of the estimated diffusion components in physical space, with an extreme corresponding to the MHB itself (Figure 3F-G; Methods). Using a spatial vector field to capture local magnitude and direction of changes in diffusion component 1 in space, we observed an outward radiation of signaling gradients from the MHB region, corresponding to the rostral-caudal axis (Figure 3G), with strong enrichment for *Lmo1*^80^ in the midbrain and *Pax8*^81^ in the hindbrain (Figure 3I). Additionally, we observed that diffusion component 2 corresponds to an emerging dorsal-ventral axis (Figure 3H), demonstrating that the coordinate space of the brain is established at this stage of development.

To identify genes contributing towards to the emergence of this coordinate space, we performed unbiased detection of spatially variable genes (Methods^82^; Supplementary Figure 20; Supplementary Table 7), uncovering distinct spatial expression patterns, especially along the dorsal-ventral axis within the hindbrain (Supplementary Figure 20). Among spatially variable genes, several are known regulators of cell fate commitment including *Fgf8, Fgf17, Wnt1*, and *En1*, all of which displayed their highest level of expression at the MHB (Figure 3I). *Fgf8* is a known MHB organizer, whose posterior expression relative to the boundary is necessary for repressing the expression of *Otx2* in the rhombencephalon^83^. Consistent with this, we inferred that the imputed expression of *Fgf8* was negatively correlated with *Otx2*. By contrast, *Wnt1*, whose imputed expression is restricted rostral of the MHB, is known to up-regulate *Otx2* expression in the midbrain^84,85^. *En1* (Engrailed 1) expression was observed across the MHB with no rostral or caudal bias^86–88^ (Figure 3I). Interestingly, in *Wnt1*^*−/−*^ embryos the expression of *En1* is absent, consistent with the importance of WNT1 signaling for *En1* expression^89,90^. This is supported by the observation that the deletion of *En1* results in a midbrain-hindbrain deletion, with a phenotype that closely resembles the *Wnt1*^*−/−*^ mutant mice^86^. We also observed spatially-distinct expression of *Foxa2* and *Shh* in the floor plate, another important midbrain organizer (Figure 3I), consistent with the observation that both genes are critical for the specification of the floor plate^91^. Additionally, we observed a cluster of cells, characterized by the highly restricted inferred expression of *Msx3*, in the dorsal developing neuronal tube^92^. Finally, we observed that *Ezr* (Ezrin), *Efna2* (Ephrin A2) and *Efnb1* (Ephrin B1) were among the genes with the most spatially variable patterns of expression. The Ephrin signaling pathway is a known regulator of cell sorting and plays an important role in the formation of a sharp MHB that compartmentalizes the brain^93^. Consistent with this, *Efna2* and *Efnb1* are inferred to occupy distinct territories of gene expression on each side of the MHB. Taken together, this analysis demonstrates how the imputed data can be used to reliably recapitulate and enhance our understanding of important developmental process, such as MHB formation. In particular, it captures the early transcriptional effects of neural tube rostro-caudal and dorso-ventral regionalization (Figure 3G-H) concomitant with the establishment of the MHB and floor plate signaling hubs.

### Spatial patterning of cells within the gut tube is associated with cell type identity

Finally, we examined the emergence of organ precursor cells along the anterior-posterior axis in the developing gut tube. Recently, Nowotschin *et al.* inferred the pseudo-spatial ordering of E8.75 (13 somite stage(ss)) gut tube cells along the anterior-posterior axis^2^. However, despite validation of the anterior-posterior patterning using targeted *in situ* hybridizations, the ability to finely determine the boundary between cell types and to precisely demarcate the locations of cell types along complex tissues like the gut tube is challenging when using single gene *in situ* stainings. To explore whether our data could shed new light on this problem, we performed a joint mapping of the seqFISH data with cells from dissected E8.75 (13ss) gut tubes that were profiled using scRNA-seq^2^ (Figure 4A; Supplementary Figure 21). Incorporating this additional scRNA-seq dataset allowed us to further refine the cellular annotations for the developing gut tube and nearby relevant cell types – in particular, it allowed us to associate cells with the organs that they would likely contribute to in the adult animal, including thyroid, thymus, lung, liver, pancreas, small intestine, large intestine/colon. Notably, the seqFISH profiled embryos, in comparison to the Nowotschin dataset, lack cells associated with the large intestine, likely due to the area of the large intestine not being represented in the tissue sections profiled using seqFISH (Supplementary Figure 21).

**Figure 4:**
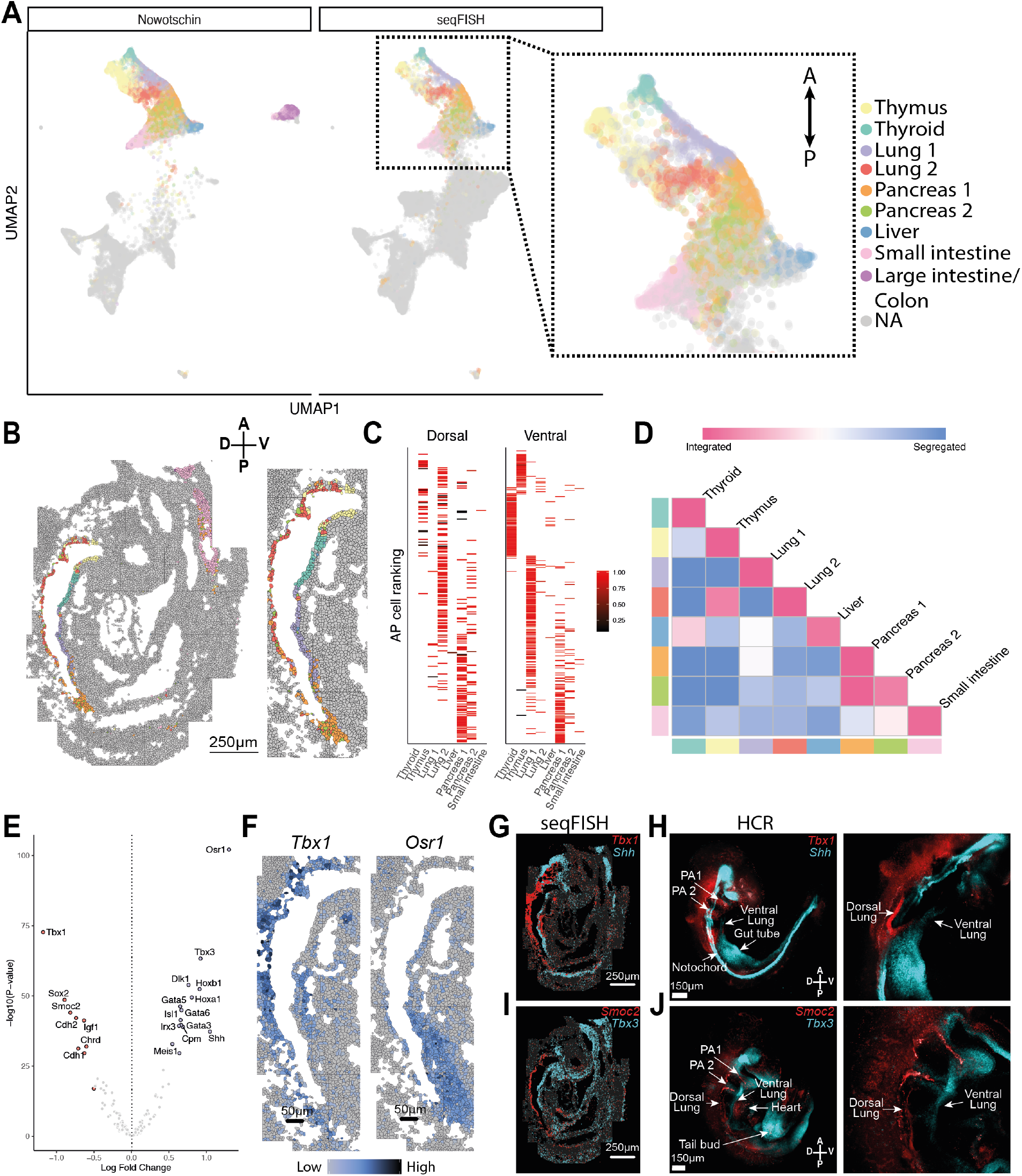
Spatial characterization of gut tube organogenesis. (A) Joint embedding of seqFISH data and Nowotschin *et al.* cells corresponding to the developing gut tube^2^ with seqFISH cells annotated by their predicted gut tube subtype. Colors represent gut tube subtypes. Zoomed in region shows anterior-posterior patterning of the gut endoderm cluster in the UMAP space, indicated by arrow. Abbreviations used: A= anterior; P= posterior. (B) Position of gut tube cell types in the embryo tissue section. Colors represent cell type classification. Scale bar 250 μm. Right hand side shows a zoom in into the region of the gut tube for better visualization. (C) Anterior-posterior ranking of cells, corresponding to each gut tube subtype, split into dorsal and ventral regions. Bar color corresponds to the mapping score associated with classification into the subtype. (D) Cell-cell contact map that displays the relative enrichment towards integration and segregation of pairs of gut tube subtypes in space, ordered along the inferred A-P ordering in Nowotschin *et al.*^2^. (E) Volcano plot showing gene expression comparison between the (ventral) Lung1 and (dorsal) Lung2 subtypes using seqFISH data. Significantly differentially expressed genes (absolute LFC > 0.5 & FDR-adjusted P-value < 0.05) are highlighted and corresponding gene names are indicated. (F) Visualization of expression of *Tbx1* (enriched in the dorsal Lung2 cluster) and *Osr1* (enriched in the ventral Lung1 cluster). Scale bar 50 μm. (G) ‘Digital *in situ*’ showing detected mRNA molecules for *Tbx1* (red) and *Shh* (cyan) across the entire embryo tissue section. Scale bar 250 μm. (H) Multiplexed mRNA imaging of whole-mount E8.75 mouse embryo using hybridization chain reaction (HCR) of *Tbx1* (red) and *Shh* (cyan). Zoom in shows region specific expression in the developing lung region. Scale bar 150 μm. Abbreviation used: PA= pharyngeal arch. (I) ‘Digital *in situ*’ showing detected mRNA molecules for *Smoc2 (red)* and *Tbx3* (cyan) across the entire embryo tissue section. Scale bar 250 μm. (J) Multiplexed mRNA imaging of whole-mount E8.75 mouse embryo using hybridization chain reaction (HCR) of *Smoc2* (red) and *Tbx3* (cyan). Zoom in shows region specific expression in the developing lung region. Scale bar 150 μm.

As expected, plotting the physical position of the subclusters showed distinct patterning along the anterior-posterior axis (Figure 4B). Interestingly, this patterning of cell types along the anterior-posterior axis was mirrored by the presence of spatially-distinct populations of cells within the surrounding splanchnic mesoderm (Methods; Supplementary Figure 22), consistent with recent reports^3^ and confirming that signalling interactions between the gut endoderm and the surrounding mesoderm plays a key role in determining cell type identity^92^.

Topological cell-cell contact analysis of all gut tube subclusters revealed a spatial separation of two lung subtypes (Lung1 and Lung2) defined by Nowotschin *et al.* (Figure 4C-D). We observed that cells assigned a Lung1 identity were located exclusively on the ventral side of the gut tube, while Lung2 cells were located on the dorsal side, suggesting an early symmetry breaking event (Figure 4B; Supplementary Figure 23). To further understand the spatial separation of the two Lung subclusters, we performed a differential gene expression analysis. As expected, we observed gene expression changes associated with dorsal-ventral patterning (Figure 4E; Supplementary Table 8), including differential expression of *Chordin*, a known dorsal-ventral regulator^94^, and *Osr1*, which is necessary for lung specification and whose loss results in significantly fewer respiratory progenitors at E9.5 and reduced lung size^95^ (Figure 4F). Additionally, the T-box gene *Tbx1*, which is known to be expressed in the embryonic mesoderm and later in the pharyngeal region and otic vesicle^96^, was more strongly expressed on the dorsal side of the gut tube^96,97^. It has been demonstrated that mutants which show esophageal atresia / trachea-esophageal atresia display abnormal expression of *Tbx1*^98^ and *Tbx2*^97^. To independently validate these asymmetric dorsal-ventral expression patterns, we used whole-mount hybridization-chain reaction (HCR), combined with 3D imaging, to study the co-expression of *Tbx1* (dorsal) and *Shh* (ventral) as well as *Smoc2* (dorsal) and *Tbx3* (ventral) (Figure 4G-J; Supplementary Figure 23). This confirmed the observations from our seqFISH data, with clear dorsal-ventral localization of these genes being observed in the foregut region of the gut tube corresponding to the Lung1 and Lung2 populations.

Taken together, the spatially-resolved expression pattern of genes involved in esophagus, lung and trachea development and the anatomical position of the Lung1 and Lung2 populations indicate that the dorsal Lung2 population corresponds to esophageal progenitors, while the ventral Lung1 population represents lung and trachea progenitors. Although little is known about the transcriptional identity of the early dorsal and ventral endodermal population that ultimately give rise to the trachea and esophagus, Kuwahara *et al.* recently used single-cell RNA sequencing at E10.5 and E11.5 to better define the transcriptional identity of the developing esophagus, trachea and lung^99^. Several of the identified markers already show dorsal-ventral asymmetries in our data, including the lung and trachea marker *Isl1, Isx2* and *Isx3* and the esophagus marker *Sox2*^99–101^. More broadly, previous studies have shown that the commitment of progenitor cells to either the lung/trachea or the esophagus is coordinated by the interplay of several transcription factors and signaling pathways that also regulate the dorsal-ventral specification of the gut tube^99^. Specifically, it was shown in E9.5 embryos that the expression of *Wnt2/2b*, *Bmp4* and *Nkx2-1* is enriched in the ventral foregut and respiratory mesenchyme, while the BMP signaling inhibitor *Noggin* and *Sox2* are enriched in the dorsal foregut^3,102–105^. Consistent with this, we observe strong expression of *Wnt2/2b* and *Bmp4* in the splanchnic mesoderm surrounding the ventral Lung1 population, indicating an early role of WNT and BMP signaling in lung and trachea instruction (Supplementary Figure 22). Taken together our data suggest that the separation of cells committed to either the lung and trachea (Lung1) or the esophagus (Lung2) is already present at the 8-12 somite stage, approximately 12-24 hours earlier than previously reported.

## Discussion

In this study we have combined cutting-edge experimental approaches with advanced computational analyses to generate a comprehensive map of how gene expression varies in space across sagittal sections of an entire mouse embryo at the 8-12 somite stage of development. Previous studies using scRNA-seq have computationally reconstructed developmental trajectories based on gene expression but, in the absence of cell-specific spatial information, it has been impossible to define how cell states are correlated with the position of cells within the embryo, or to understand how the local signaling environment to which they are exposed might impact their molecular signature and their ultimate fate. Conversely, although pioneering studies have mapped the expression of individual developmental genes at single-cell resolution, the ability to stitch together multiple independent *in situ* maps into a complete, single-cell resolution map has not been possible due to inevitable fine scale variations in local cellular organization between embryos.

By combining our high-resolution seqFISH map with scRNA-seq we have delineated the precise location of distinct cell types within a single reference scaffold. To illustrate the potential of this resource, we have shown how it can provide insight into the formation of the midbrain-hindbrain boundary and, in particular, the aetiology of cell types along the nascent gut tube. In the latter case, we have added an additional axis of resolution to previous studies by uncovering dorsal-ventral patterning associated with the commitment of cells towards either the esophagus or the lung and trachea. To enable this analysis, we developed computational tools for probe design, for integrating and imputing data, as well as strategies for downstream analysis, including modelling spatial heterogeneity and for performing virtual dissections. This provides a robust experimental and computational framework for future studies, both in the mouse and in other biological systems.

In the future, the generation of comprehensive cell-resolution spatial maps at additional stages of mouse development will allow spatiotemporal analysis and provide insight into the complex processes associated with cell fate specification during gastrulation and organogenesis. 3D wholemount maps would further resolve the processes associated with embryo patterning, in particular processes that are associated with the left-right axis. Moreover, the recent development of novel image-based cell lineage tracing methods, such as Zombie^106^ or intMEMOIR^107^, allow a cell’s lineage to be recorded while preserving spatial information. These methods are compatible with seqFISH and therefore afford the possibility to record spatial gene expression profiles and cell history from the same cell in intact tissue. Combining these novel lineage tracing methods with spatial transcriptomics will improve our ability to decipher the mechanisms underpinning cell fate choice and tissue patterning.

## METHODS

### Library design

We selected genes whose expression patterns discriminated cells from different labeled cell types described in the scRNA-seq data of Pijuan-Sala *et al*. 2019^6^. To do this, we used the scran function findMarkers^108^, with the option ‘pval.type = “any”’, testing against an absolute fold change of 0.5. This was performed separately at each developmental stage of the Gastrulation atlas (E6.5-E8.5, in 0.25-day steps), and only cell types with more than 10 cells at any given stage were included in the per stage analysis. Genes were excluded if the upper quartile of the normalized count across cells in any individual cell type was greater than 20. This was performed to prevent the inclusion of highly expressed genes that may compromise imaging. The “top” 5 genes per cell type were saved from each stage, and the union of these genes was taken across stages. “Top” genes were defined by the findMarkers `Top` column, which identifies a minimal number of genes required to separate any cell type from any other. The gene panel was evaluated on a per-gene basis to exclude any genes that were too short or repetitive to produce reliable fluorescence *in situ* hybridization (FISH) probes. Additionally, for each cell type, the panel of genes was manually curated in order to ensure that the total normalized RNA count across cells for each cell type was less than 300 (Supplementary Figure 1). Finally, after determining a suitable set of cell type marker genes, we manually added genes of interest (especially transcription factors) to the panel, and iteratively performed the “fluorescent load” testing and gene removal as described in the previous two sentences. In total we selected 387 genes, of which 351 genes were detected using seqFISH and 36 using non-barcoded sequential single molecular FISH (smFISH) imaging.

### Primary probe design

Gene specific primary probes were designed for the selected 351 seqFISH and 36 smFISH genes, as previously introduced by Eng *et al.*^109^ (Supplementary Table 1). To design 30-nucleotide primary probe sequences for the 351 selected seqFISH and 36 smFISH genes, we extracted 30-nucleotide sequences of each of the selected genes, using the coding region of each gene. The mask genome and annotation from the University of Santa Cruz (UCSC) were used to look up the gene sequences. All probe sequences were selected to have a GC content in the range from 45 to 65% and to not have five or more consecutive bases. Genes with more than 48 primary probes were used as a secondary filter to remove off targets. Any gene that did not achieve a minimum of 28 probes for seqFISH and 17 probes for smFISH was dropped. To validate the specificity of the generated primary probes and to minimize off targets, we performed a BLAST search against the mouse transcriptome and all BLAST hits other than the target gene with a 15-nucleotide match were considered off targets. To avoid off target hits between the primary probes a second round of optimization was performed. We constructed a local BLAST database from the primary probe sequences and probes that were predicted to hit more than 7 times by all of the combined primary probes in the BLAST database were iteratively dropped from the probe set, until no more than 7 off-targets hits existed for each primary probe sequence.

### Readout probe design

Readout probes of 15-nucleotide length were designed as previously introduced by Shah *et al*. ^27^. In brief, the probe sequences were randomly generated with combinations of A, T, G or C nucleotides, with a GC-content in the range of 40-60%. To validate the specificity of the generated readout sequences, we performed a BLAST search against the mouse transcriptome. To minimize cross-hybridization of the readout probes, all probes with ten contiguously matching sequences between the readout probes were removed. The reverse complements of these readout-probe sequences were included in the primary probe, as described below (Primary probe library construction; Supplementary Table 1).

### Primary probe library construction

The primary probe library, consisting of 15,989 probes for 387 genes (17-48 per gene / average of 41.32 per gene), was ordered as an oligoarray pool from Twist Bioscience. Each probe for barcoded mRNA seqFISH was assembled out of 30-nucleotide mRNA complementary sequence for *in situ* hybridization, four 15-nucleotide gene specific readout sequences separated by 2-nucleotide spacer, and two flanking primer sequences to allow PCR amplification of the probe library (Primary barcoded mRNA seqFISH probes: 5’ – [primer 1] – [readout 1] – [readout 2] – [probe] – [readout 3] – [readout 4] – [primer 2] – 3’). Each of the probes for non-barcoded sequential smFISH were assembled in the same way, with the exception that the sequence for the four readout sequences was the same for all four readout sequences (Primary non-barcoded sequential smFISH probes: 5’ – [primer 1] – [readout 1] – [readout 1] – [probe] – [readout 1] – [readout 1] – [primer 2] – 3’). We used validated primer and 84 readout sequences, previously used in seqFISH+ ^26^. Next, the probe library was amplified as previously described^25,26,109–111^. In brief, limited cycle PCR was used to generate *in vitro* transcription template, using primer 1 and primer 2. Next, the PCR product was purified using a QIAquick PCR Purification Kit (Qiagen, 28104), following the manufacturer’s instructions. Subsequently, the purified PCR product was used for *in vitro* transcription (NEB, E2040S) followed by reverse transcription (Thermo Fisher, EP7051) with the forward primer containing a uracil nucleotide^112^. Next, the forward primer sequence was removed by cleaving off the uracil nucleotide. The probes were subjected to a 1:30 dilution of uracil-specific excision reagent (USER) enzyme (NEB, N5505S) for about 24h at 37 °C. The single-stranded DNA (ssDNA) was alkaline hydrolyzed with 1 M NaOH at 65 °C for 15 minutes, followed by neutralization with 1 M acetic acid to remove the remaining RNA templates. Next, the probe library was purified by ethanol precipitation to remove residual nucleotides and by phenol-chloroform extraction to remove the proteins. Finally, the amplified primary probe library was dried by speedvac and resuspended at a concentration of 40 nM per probe in primary probe hybridization buffer, composed of 40% formamide (Sigma, F9027), 2x SSC, and 10% (w/v) dextran sulphate (Sigma, D8906). The probes were stored at −20 °C.

### Readout probe synthesis

15-nucleotide readout probes were ordered from Integrated DNA Technologies (IDT), conjugated to Alexa Fluor 488, Cy3B and Alexa Fluor 647 as indicated in Supplementary Table 2 and 3. All readout probes were stored at −20 °C.

### Encoding strategy

In this experiment we used a 12-pseudocolour encoding scheme, as described previously^27,109^. In brief, 12-pseudocolours were equally separated across three fluorescent channels (Alexa Fluor 488, Cy3B and Alexa Fluor 647). The 12-pseudocolour imaging was repeated four times, resulting in 12^4^ (20,736) unique barcodes. Additionally, an extra round of pseudocolour imaging was performed to obtain error-correctable barcodes, as previously introduced^25^. In this experiment, 351 genes were encoded across all channels (Supplementary Table 2).

### Coverslip functionalization

Coverslips were functionalized as previously described ^26^. In brief, coverslips (Thermo Scientific, 3421) were washed in nuclease free water, followed by an immersion in 100% ethanol (Koptec). Subsequently, coverslips were air dried and cleaned using a plasma cleaner on the high setting (PDC-001, Harrick Plasma) for 5 minutes. Then, the coverslips were immersed in 1% bind-saline solution (GE, 17-1330-13) made in pH 3.5 10% (v/v) acidic ethanol solution for 1 hour at room temperature. Next, coverslips were rinsed three times in 100% ethanol and heat-dried in an oven at >90 °C for 30 min. Then, the coverslips were treated with 100 μg/ml of Poly-D-lysin (Sigma, P6407) in water for a minimum of 1 hour at RT. Afterwards, coverslips were washed three times in nuclease free water and air dried. Functionalized coverslips can be stored for up to 1-week at 4 °C.

### Mice

Experiments, with exception of the HCR experiment (see below), were performed in accordance with EU guidelines for the care and use of laboratory animals, and under authority of appropriate UK governmental legislation. 8-12 week wild-type C57BL/6J mice (Charles Rivers) were used, with exception of the HCR experiment (see below). For the HCR experiment, wild-type CD-1 mice (Charles Rivers) were used. Mice used were housed under a 12-h light/dark cycle. Natural mating was set up between males and 4–6-week-old virgin females, with noon of the day of vaginal plug considered to be E0.5. Mice were maintained in accordance with guidelines from Memorial Sloan Kettering Cancer Center (MSKCC) Institutional Animal Care and Use Committee (IACUC) under protocol no. 03-12-017 (principal investigator A.-K.H.).

### Tissue preparation

Embryos were dissected from the uteri, washed in M2 media (Sigma Aldrich, 7167) and immediately placed in 4% PFA (Thermo Scientific, 28908) in 1x PBS (Invitrogen, AM9624) for 30 minutes at room temperature. The embryos were then washed and immersed in 30 % RNase-free sucrose (Sigma Aldrich, 84097) in 1x PBS at 4 °C until the embryo sank to the bottom of the tube. Afterwards, each embryo was positioned in a sagittal orientation in a tissue base mold (Sakura, 4162) in optimal cutting temperature compound (OCT) solution (Sakura, 4583) and frozen in a dry ice isopropanol (VWR, 20842) and stored at −80 °C. 20 μm tissue section were cut using a cryotome and collected on the functionalized coverslips and stored at −80 °C.

### seqFISH using tissue sections

Tissue sections were post-fixed with 4% PFA in 1x PBS for 15 minutes at room temperature to stabilize the DNA, RNA and overall sample structure. The fixed samples were permeabilized with 70% EtOH for 1 hour at room temperature. Then the tissue slices were cleared with 8% SDS in 1x PBS for 20 minutes at room temperature. The cleared sample was washed with 70% EtOH and then air-dried. Samples were blocked for a minimum of 2 hours in blocking solution at room temperature in a humidified chamber (1x PBS, supplemented with 0.25% TritionX-100, 10 mg/ml BSA (Thermo Fisher, AM2616), 0.5 mg/ml salmon sperm DNA (Thermo Fisher, AM9680)). Anti-pan Cadherin (Abcam, ab22744), anti-N-Cadherin (Cell Signaling Technology, [13A9], 14215), anti-β-Catenin antibody (15B8) (Abcam, ab6301), and anti-E-Cadherin antibody (BD Biosciences, clone 36, 610181) were diluted in blocking solution and incubated for 2 hours at room temperature. Samples were washed three times in 1x PBS, supplemented with 0.1% TritonX-100 (PBS-T), before incubating anti-mouse IgG secondary antibody, conjugated to CCTTACACCAACCCT oligo, diluted 1:500 in blocking solution for at least 2 hours at room temperature. Next, the samples were washed three times in 1x PBS-T. The samples were post-fixed with 4% PFA in 1x PBS for 15 minutes followed by three 10-minute washes in 2x SSC (Thermo Fisher, 15557036). The samples were dried and hybridized for 24-36 hours with the probe library (~2.5 nM per probe), 1 nM of *Eef2* probe set A and B (Supplementary Table 1), and 1 μM Locked Nucleic Acid (LNA) oligo-d(T)30 (Qiagen) in primary-probe hybridization buffer composed of 40% formamide (Sigma, F9027), 2x SSC and 10% (w/v) dextran sulfate (Sigma, D8906) in a humid chamber at 37 °C. The hybridization samples were washed with 40% formamide wash buffer (40% formamide, 0.1% TritonX-100 in 2x SSC) for 30 minutes at 37 °C, followed by three rinses with 2x SSC. Then, the samples were hybridized for at least 2 hours with 200 nM tertiary probe (/5Acryd/AG GGT TGG TGT AAG GTT TAC CTG GCG TTG CGA CGA CTA A) in EC buffer made of 10% ethylene carbonate (Sigma, E26258), 10 % dextran sulfate (Sigma, D4911), 4x SSC. The samples were washed for 5 minutes in a 10% formamide washing buffer (10% formamide, 0.1% TritonX-100 in 2x SSC), followed by two 5-minute washes in 2x SSC. The samples were treated with 0.1 mg/ml Acryoloyl-X succinimidyl ester (Thermo Fisher, A20770) in 1x PBS for 30 minutes at room temperature. Then the samples were rinsed three times with 2x SSC and post-fixed with 4% PFA in 1x PBS for 15 minutes, followed by three washes in 2x SSC. Next, the samples were incubated with 4% acrylamide/bis (1:19 crosslinking) hydrogel solution in 2x SSC for 30 minutes. The hydrogel solution was aspirated and the sample covered with 20 μl of degassed 4% hydrogel solution containing 0.05% ammonium persulfate (APS) (Sigma, A3078) and 0.05% *N,N,N’,N’-*tetramethylenediamine (TEMED) (Sigma, T7024) in 2x SSC. The sample was sandwiched by GelSlick functionalized slide (Lonza, 50640). The samples were transferred to a home-made nitrogen gas chamber and incubated for 30 minutes at room temperature, before transferring to 37 °C for at least 3 hours. After polymerization, the slides were gently separated from the coverslip and the hydrogel-embedded tissue was rinsed with 2x SSC three times. Then the samples were cleared for 3 hours at 37°C using digestion buffer, as previously described^34^. The digestion buffer consisted of 1:100 proteinase K (NEB, P8107S), 50 mM pH 8 Tris-HCl (Invitrogen, AM9856), 1 mM EDTA (Invitrogen, 15575020), 0.5% Triton-X100, 1% SDS and 500 mM NaCl (Sigma, S5150). After digestion, the tissue slices were rinsed with 2x SSC multiple times and then subjected to 0.1 mg/ml label-X modification for 45 minutes at 37°C^34^. For further stabilization the sample was re-embedded in a 4% hydrogel solution as described above, with a shortened gelation time of 2.5 hours. Excess gel was removed with a razor and the sample covered with an in-house made flow cell. The sample was immediately imaged.

### seqFISH imaging

Two tissue sections from two experimental blocks, containing three embryos, were imaged as previously described^26,27^. In brief, the flow cell was connected to an automated fluidics system. First, the sample was stained with 10 μg/ml DAPI (Sigma, D8417) in 4x SSC and the field of view (FOV) were selected. All rounds of imaging were performed in anti-bleaching buffer made of 50 mM Tris-HCl pH 8.0 (Thermo Fisher, 15568025), 300 mM NaCl (Sigma, S5150), 2x SSC (Thermo Fisher, 15557036), 3 mM Trolox (Sigma, 238813), 0.8% D-glucose (Sigma, G7528), 1:100 diluted Catalase (Sigma, C3155), and 0.5 mg/mL Glucose oxidase (Sigma, G2133). The RNA integrity of the sample was validated by colocalization of the dots of two interspersed *Eef2* probes, each read out by secondary readout probes with distinct fluorophores (Supplementary Figure 2; Supplementary Table 3). Sixteen hybridization rounds were imaged for the decoding of the barcoded mRNA seqFISH probes followed by a repeat of the first hybridization. Then, twelve-serial hybridization rounds were imaged for 36 non-barcoded sequential smFISH probes, followed by one hybridization to visualize the cell segmentation staining, using Cy3B conjugated to /5AmMC6/TTAGTCGTCGCAACG. The hybridization buffer for each of the hybridization rounds, excluding the last, contained three unique readout probes, each consisting of a unique 15 nucleotide probe sequences, conjugated to either Alexa Fluor 647 (50 nM), Cy3B (50 nM) or Alexa Fluor 488 (50 nM) in EC buffer, as described above (Supplementary Table 2-3). The hybridization buffer for the cell segmentation staining contained one unique 15-nucleotide probe sequence conjugated to Alexa Fluor 647. The hybridization buffer mixes for the 30 rounds of hybridization were stored in a deep bottom 96-well plate and were added to the hybridization chamber by an automated sampler system, as described previously^26^. The tissue section was incubated in the hybridization solution for 25 minutes at room temperature in the dark. Next, the sample was washed with 300 μl of 10% formamide wash buffer to remove excess and non-specific readout probes. The sample was rinsed with 4x SSC and subsequently stained with 10 μg/ml DAPI in 4x SSC for 1.5 minutes. Then, the flow chamber was filled with anti-bleaching buffer and all selected FOVs of the sample were imaged. The microscope used was a Leica DMi8 stand equipped with a Yokogawa CSU-W1 spinning disk confocal scanner, an Andor Zyla 4.2 Plus sCMOS camera, a 63x Leica 1.40 NA oil objective, a motorized stage (ASI MS2000), lasers from CNI and filter sets from Semrock. For each field of view, snapshots were acquired with 4 μm z-steps for 6 z-slices. After imaging, the readout probes were stripped off using 55% wash buffer (55% formamide, 0.1% Triton-X100 in 2x SSC) by incubating the sample for 4 minutes, followed by 4x SSC rinse. Serial hybridization and imaging were repeated for 29 rounds. The integration of automated fluidics delivery system and imaging was controlled by a custom script written in μManager^113^.

### Image processing

To remove the effects of chromatic aberration, 0.1 mm TetraSpeck beads’ (Thermo Scientific T7279) images were first used to create geometric transforms to align all fluorescence channels. Tissue background and auto-fluorescence were then removed by dividing the initial background with the fluorescence images. To correct for the non-uniform background, a flat field correction was applied by dividing the normalized background illumination with each of the fluorescence images while preserving the intensity profile of the fluorescent points. The background signal was then subtracted using the ImageJ rolling ball background subtraction algorithm with a radius of 3 pixels and filtered with a despeckle algorithm to remove any hot pixels.

### Image registration

Each round of imaging contained the 405 channel, which included the DAPI stain of the cell. For each field of view (e.g. tile), all of the DAPI images from every round of hybridization were aligned to the first image using a 2D phase correlation algorithm.

### Cell segmentation

For semi-automatic cell segmentation, the membrane stains β-catenin, E-cadherin, N-Cadherin and Pen-cadherin were aligned to the first hybridization round using DAPI, and subsequently trained with Ilastik^36^, an interactive supervised machine learning toolkit, to output probability maps, which were used in the Multicut^114^ tool to produce 2D labeled cells for each z-slice. For image analysis, potential mRNA transcript signals were located by finding the local maxima in the processed image above a predetermined pixel threshold, manually calculated for one field of view and adjusted for the remainder according to the number of expected potential spots per cell. The transcript spots were assigned to the corresponding labeled cells according to location, thereby generating a gene-cell count table.

### Barcode calling

Once all potential points in all channels of all hybridizations were obtained, dots were matched to potential barcode partners in all other channels of all other hybridizations using a 2.45-pixel search radius to find symmetric nearest neighbors. Point combinations that yielded only a single barcode were immediately matched to the on-target barcode set. For points that matched to multiple barcodes, first the point sets were filtered by calculating the residual spatial distance of each potential barcode point set and only the point sets giving the minimum residuals were used to match to a barcode. If multiple barcodes were still possible, the point was matched to its closest on-target barcode with a hamming distance of 1. If multiple on target barcodes were still possible, then the point was dropped from the analysis as an ambiguous barcode. This procedure was repeated using each hybridization as a seed for barcode finding and only barcodes that were called similarly in at least 3 out of 4 rounds were validated as genes. For more details regarding the seqFISH method, please refer to Shah *et al*.^25^.

### smFISH processing

For the 36 genes that were probed using smFISH, twelve sequential rounds of imaging across three fluorescent channels (corresponding to A647, Cy3B and A488 respectively) were used (Supplementary Table 3). Assignment of an optimal light intensity threshold to separate background noise from hybridized mRNA molecules poses an additional challenge for these data since, unlike the seqFISH probed transcripts, each gene’s expression is measured only over a single round of hybridization.

To address this problem, we manually assigned a threshold for three randomly selected fields of view in the first experimental block (corresponding to embryos 1 and 2) and three fields of view in the second experimental block (embryo 3) for all fluorescent channels and all hybridization rounds. The choice of threshold was motivated by considering the minimum value at which we acquire nearly complete loss of dots in cell-free areas, which we expect should only contain background signal. We then assessed the relationship between the channel, and hybridization round and the manually selected thresholds, observing that intensity thresholds are highly channel specific, but do not vary as a function of hybridization round (Supplementary Figure 24). Accordingly, for each channel, hybridization round and experimental block, we assigned the intensity threshold as the average across all manually selected thresholds.

We then visually assessed the spatial distribution of selected spots for each gene, embryo and z-slice. While most of estimated intensity thresholds resulted in spatially coherent expression patterns across all embryos, we noticed a strong channel - field of view specific effect for some genes. Specifically, in the first experimental block, genes probed with A647 exhibited substantial background signal in fields of view 39, 40 and 44. Given that the effect is highly specific to this channel, it is likely an artefact of the imaging experiment. For these genes and fields of view, manual examination of a wide range of appropriate intensity thresholds failed to identify a threshold at which the background noise was eliminated (Supplementary Figure 24). Consequently, we discarded these fields when evaluating the performance of our imputation strategy (see below).

### Whole-mount hybridization chain reaction (HCR) on E8.75 mouse embryos

Hybridization chain reaction fluorescent *in situs* where carried out as described^115,116^ with the modification of using 60 pmol of each hairpin per 0.5ml of amplification buffer. Hairpins were left 12-14 hours at room temperature for saturation of amplification to achieve highest levels of signal to noise^117^. Split initiator probes (V3.0) were designed by Molecular Instruments, Inc.

### HCR imaging

All images were obtained on a Zeiss 880 laser scanning confocal microscope with a 10x objective and 6.74 μm z-step sizes. Tile-scanned z-stacks were stitched in Zen software and rendered in 3D in Imaris (v9.6, Bitplane Inc).

### Downstream computational analysis

#### Quality control and filtering

To lower the chance of counting cells multiple times in contiguous z-slices, we selected two z-slices (denoted 1 and 2 hereafter) for further analysis, corresponding to two parallel tissue layers 12um apart. We then removed segmented regions most likely to correspond to empty space rather than cell-containing regions by testing whether a putative cell’s square-root transformed segmented area was larger than expected (Z-test; FDR threshold of 0.01). Of the remaining segmented regions, we considered segments containing at least 10 detected mRNA molecules corresponding to at least 5 unique genes as true cells.

#### Cell neighborhood network construction

To construct a cell neighborhood network, for each cell within a given embryo and z-slice we extracted the polygon representation of the cell’s segmentation, corresponding to a set of vertex coordinates. We then calculated an expanded segmentation by constructing a new polygon where each expanded vertex was lengthened along the line containing the original vertex and the center of the polygon. We performed a multiplicative expansion of 1.3 for each vertex. To construct the cell neighborhood network, we then identified the other cells in which segmentation vertices were found to be within the expanded polygon. Cell neighborhood networks were considered separately for each embryo and z-slice combination.

#### Gene expression quantification per cell

We calculated normalized expression logcounts for each cell using scran’s logNormCounts function^108^, with size factors corresponding to the total number of mRNAs (excluding the sex-specific gene *Xist*) identified for each cell. Size factors were scaled to unity and a pseudocount of 1 was added before the logcounts were extracted. For the majority of downstream analyses, such as differential gene expression, we specifically included biological and technical variables (i.e. z-slice and field of view) as covariates. However, for the task of harmoniously visualizing gene expression in spatial coordinates, we extracted ‘batch-corrected expression’ values for each gene. This was done by first performing batch correction using the Mutual Nearest Neighbors method, implemented with fastMNN in the scran package^108^, with batch variables corresponding to z-slice and field of view. To ensure interpretability of the reconstructed expression values, we rescaled these values to correspond to the unnormalized logcounts expression distribution for each gene, resulting in a “batch-corrected expression” matrix.

#### Clustering gene expression

To identify unsupervised clusters, we first performed multi-batch aware PCA on the normalized logcounts using the multiBatchPCA function in scran^108^, with z-slice and field of view as batch variables, using all genes except *Xist* as input to extract 50 PCs. We then performed batch correction using the Mutual Nearest Neighbors approach, resulting in a corrected reduced dimension embedding of cells. To identify clusters, we estimated a shared nearest neighbor network, followed by Louvain network clustering. To further extract unsupervised subclusters, for each set of cells belonging to a given cluster we performed highly variable gene selection to select genes with a non-zero estimated biological variance, excluding the sex-specific gene *Xist*. Using these selected genes, we performed batch-aware PCA to extract 50 PCs, followed by batch correction, shared nearest neighbor network construction and Louvain clustering similar to what was performed for all cells.

#### Joint analysis with Gastrulation atlas

We downloaded the E8.5 Pijuan-Sala *et al*.^6^ 10X Genomics scRNA-seq dataset from the Bioconductor package *MouseGastrulationData* and performed batch aware normalization using the multiBatchNorm function in the scran package^108^, before extracting cells that correspond to a known cell type with at least 25 cells. Cell types associated with the somitic and paraxial mesoderm were further refined using labels assigned by Carolina Guibentif (*personal communication*); blood subtypes (Erythroid1/2/3 and Blood progenitors 1/2) were collapsed to the two major groups; ExE mesoderm was renamed to Lateral plate mesoderm; and Pharyngeal mesoderm was renamed to Splanchnic mesoderm. Subsequently, only genes probed by both the scRNA-seq and seqFISH assays were kept for this analysis. We then jointly embedded the normalized logcounts of each of the two datasets by performing batch-aware PCA with 50 components (excluding the sex-specific gene *Xist*) via the multiBatchPCA function in scran with batch variables corresponding to sequencing runs in the Gastrulation atlas and field of view and z-slice for the seqFISH data. We corrected for platform and batch specific effects using the MNN method (fastMNN^118^), ensuring that merge ordering is such that Gastrulation atlas batches are merged first (ordered by decreasing number of cells). This joint embedding of the Gastrulation atlas and seqFISH dataset was further reduced in dimension using UMAP, implemented by calculate UMAP in scran^108^ to allow the data to be visualized in two dimensions.

#### Cell type identification

To assign a cell type label to each seqFISH cell, we considered the Gastrulation atlas cells that it was closest to in the batch-corrected space. We considered the k-nearest cells, with the distance from the seqFISH cell to its Gastrulation atlas neighbors being computed as the Euclidean distance amongst the batch-corrected PC coordinates. We set the number of nearest neighbors, k, to 25. Ties were broken by favoring the cell type of those closest in distance to the query cell. We calculated a “mapping score” for each query cell as the proportion of the majority cell type present among the 25 closest cells.

To further refine the predicted cell types we performed joint clustering of the Gastrulation atlas and seqFISH cells by building a shared nearest neighbor network on the joint PCs followed by Louvain network clustering. Additionally, we further subclustered the output by building a shared nearest neighbor network on the cells corresponding to each cluster followed by Louvain network clustering. We then inspected the relative contribution of cells to each subcluster as well as the expression of marker genes in order to identify subclusters that potentially required manual re-annotation, either due to small differences in composition in the reference atlas or in the gene expression profile (Supplementary Figure 6). We also identified a set of subclusters that were likely associated with low quality cells, defined by lower total mRNA counts. Furthermore, we performed virtual dissection on regions corresponding anatomically to the developing gut tube, and for these cells re-classified those that were “Surface ectoderm” as “Gut tube”.

#### Subclustering of mixed mesenchymal mesoderm cells

To analyze the mixed mesenchymal mesoderm, population we performed highly variable gene selection for these cells only, using the ‘modelGeneVar’ function in scran^108^, and performed principal component analysis (excluding the sex-specific gene *Xist*) on the normalized logcounts followed by batch correction using MNN with embryo and z-slice as batch variables. We then further reduced these corrected PCs into two dimensions using UMAP for visualization purposes. To identify mixed mesenchymal mesoderm subclusters, we estimated a shared nearest neighbor network, followed by Louvain network clustering. We then performed differential expression analysis on the seqFISH genes and on the imputed gene expression values (described further below) using the ‘findMarkers’ function in scran^108^, and gene ontology enrichment analysis as described below. To further identify the spatial context for the mixed mesenchymal mesoderm, for each cluster we extracted the cells that appear as direct contact neighbors with any cell belonging to the cluster, and recorded their corresponding cell type. To assess the relative association of each mixed mesenchymal mesoderm subcluster to the Gastrulation atlas^6^, we calculated a weighted score per Gastrulation atlas cell and mixed mesenchymal mesoderm subcluster, corresponding to the average ranking of the Gastrulation atlas cell among the top 25 nearest neighbors for each mixed mesenchymal mesoderm subcluster cell.

#### Spatial heterogeneity testing per cell type

We identified genes with a spatially heterogeneous pattern of expression using a linear model with observations corresponding to each cell for a given cell type, and with outcome corresponding to the gene of interest’s expression value. For each gene, we fit a linear model including the embryo and z-slice information as covariates as well as an additional covariate corresponding to the weighted mean of all other cells’ gene expression values. The weight was computed as the inverse of the cell-cell distance in the cell-cell neighborhood network.

More formally, let x_ij_ be the expression of gene *i* for cell *j*. Calculate 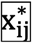 as the weighted average of other *K* cells’ expression, weighted by distance in the neighborhood network

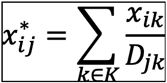

where

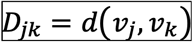

is the path length in the neighborhood network between vertices corresponding to cells *j* and *k*.

We then fit the linear model for each gene

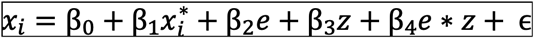

where *e* and *z* correspond to the embryo and z-slice identity of the cells, and *∈* represents random normally distributed noise. The *t*-statistic corresponding to *β*_1_ is reported here as a measure of spatial heterogeneity for the given gene, a large value corresponding to a strong spatial expression pattern of the gene in space, especially among its neighbors.

#### Subclustering of developing brain cells

To further subcluster the developing brain cells, we extracted the Gastrulation atlas cells corresponding to embryonic day E8.5 that were classified as Forebrain/Midbrain/Hindbrain. For these cells we identified genes to further cluster by using the scran function modelGeneVar^108^ to identify highly variable genes with nonzero biological variability, excluding the sex-specific gene *Xist*. For these genes we extracted the cosine-standardized logcounts, which were standardized against the entire transcriptome. We then performed batch correction using the MNN method on batch-aware PC coordinates, where batches corresponded to the sequencing samples. Using this batch-corrected embedding we estimated a shared nearest neighborhood network and performed Louvain network clustering. To relate these brain subcluster labels to the seqFISH data, we extracted the nearest neighbor information (as described in “Cell type identification”) for seqFISH cells corresponding to Forebrain/Midbrain/Hindbrain, and classified their brain subcluster label using k-Nearest Neighbors with k = 25, with closest cells breaking ties. We then named these subclusters based on marker gene expression, including a class that may be technically driven (NA class).

#### Cell-cell contact map inference

We constructed cell-cell contact maps for multiple cell annotation labelings, including mapped cell types, subclusters within each cell type, and for mapped gut tube subtypes. To do this, for each embryo and z-slice combination, we extracted the cell neighborhood network and cell-level annotation. We then generated cell-cell contact maps by first calculating the number of edges for which a particular pair of annotated groups was observed. We then randomly re-assigned (500 times) the annotation by sampling without replacement, and calculated the number of edges for all pairs of annotated groups. To construct the cell-cell contact map, we reported the proportion of times the randomly re-assigned number of edges was larger than or equal to the observed number of edges. Small values correspond to the pair of annotation groups being more segregated, and large values correspond to them being more integrated in physical space compared to a random allocation. To combine these cell-cell contact maps for each embryo and z-slice combination, we further calculated the element-wise mean for each pair of cell labels. We visualized this in a heatmap, ordering the annotation groups using hierarchical clustering with Euclidean distance and complete linkage. In the case of the gut tube subtypes, we ordered these classes by the anterior-posterior ordering given by Nowotschin *et al.*^2^. In the brain subtypes, we ordered these classes by their approximate anatomical location, from the forebrain to the hindbrain region.

#### Gene ontology enrichment analysis

To functionally annotate sets of gene clusters, we performed gene set enrichment analysis using mouse Gene Ontology terms with between 10 and 500 genes appearing in each dataset, and at least one gene appearing from the testing scaffold^119^ using Fisher’s exact test to test for overrepresentation of genes, using all scHOT tested genes as the gene universe. An FDR adjusted P < 0.05 was considered to be statistically significant.

### Imputation

Below we outline the different elements of our strategy for imputing the spatially-resolved expression of genes not profiled using seqFISH.

#### Intermediate mapping

First, for each gene in the seqFISH library (excluding the sex-specific gene *Xist*), we performed an intermediate mapping to align each seqFISH cell with the most similar set of cells in the scRNA-seq dataset. To perform the mapping we excluded the gene of interest and used the remaining 349 genes (351 seqFISH genes – *Xist* – gene of interest), resulting in 350 intermediate mappings overall. The leave-one-gene-out mapping approach was used to assess whether the intermediate mapping strategy outlined below could be used to estimate the expression counts of the omitted gene.

Similar to the integration strategy described earlier for assigning cell type labels, for each embryo and z-slice we concatenated the cosine normalized seqFISH counts with the cosine normalized expression values from the Gastrulation atlas scRNA-seq data^6^. We performed dimensionality reduction using ‘multibatchPCA’ (using 50 principal components) and performed batch correction using the ‘reducedMNN’ function implemented in scran^108^. Next, for each cell in the seqFISH dataset that was assigned a cell type identity in the earlier integration, we used the ‘queryKNN’ function in BiocNeighbors to identify its 25 nearest neighbors in the scRNA-seq data. Finally, for each seqFISH cell, the expression count of the gene of interest is estimated as the average expression of the corresponding gene across the cell’s 25 nearest neighbors.

#### Performance evaluation

For each mapped gene, its Performance score is calculated as the Pearson correlation (across cells) between its imputed values and its measured seqFISH expression level. To estimate an upper bound on the performance score (i.e., the maximum correlation we might expect to observe) we took advantage of the four independent batches of E8.5 cells that were processed in the scRNA-seq Gastrulation atlas. In particular, we treated one of the four batches as the query set and used the leave-one-out approach described above to impute the expression of genes of interest by mapping cells onto a reference composed of the remaining three batches. Additionally, to mimic the seqFISH imputation, we considered a subset of the Gastrulation atlas data consisting of only those genes that were probed in the seqFISH experiment. Moreover, due to the experimental procedure, some cell types present in the Gastrulation atlas (e.g., extra-embryonic cell types) were not probed in the seqFISH experiment. Accordingly, we considered only the subset of scRNA-seq profiled cells that were amongst the nearest neighbors of a seqFISH mapped cells so this subset most faithfully resembled the seqFISH data.

Subsequently, for each mapped gene, we computed its Prediction score as the weighted Pearson correlation between its imputed expression level and its true expression level. The weights were proportional to the number of times each Gastrulation atlas cell was present among the neighbors of a seqFISH cell, across all profiled genes.

Finally, for each gene probed in the seqFISH experiment, we define its normalized imputation performance score as the ratio of its performance score over its prediction score.

#### Final imputation

To perform imputation for all genes, we aggregated across the 350 intermediate mappings generated from each gene probed using seqFISH. Specifically, for each seqFISH cell, we considered the set of all Gastrulation atlas cells that were associated with it in any intermediate mapping. Subsequently, for every cell, we calculated each gene’s imputed expression level as the weighted average of the gene’s expression across the associated set of Gastrulation atlas cells, where weights were proportional to the number of times each Gastrulation atlas cell was present.

### Midbrain-Hindbrain Boundary (MHB) detection and virtual dissection

To identify the MHB, we visually assessed the expression of the well-described mesencephalon and prosencephalon marker *Otx2* and the rhombencephalon marker *Gbx2* (Supplementary Figure 19). We manually selected the physical region where both genes are expressed and defined this as the field of view (black rectangle, Supplementary Figure 19). Subsequently, within the selected region we performed a virtual dissection by manually choosing the boundary that best discriminates the expression of *Otx2* and *Gbx2* (Supplementary Figure 19) and based on the boundary we assigned cells either a Midbrain or a Hindbrain identity.

### Downstream analysis of the MHB region

Differential expression analysis was performed between Midbrain and Hindbrain assigned cells using the scran function ‘findMarkers’ (with a log fold-change threshold of 0.2 and an FDR-adjusted P-value threshold of 0.05; Supplementary Table 6).

To perform diffusion analysis of the MHB region, we performed batch correction of the fields of view and z-slice using the MNN approach, with logcounts of all genes excluding the sex-specific gene *Xist* as input. We then used the diffusion pseudotime (DPT) method implemented in the R package destiny^79^ to build a diffusion map with 20 diffusion components (DC), using the cell with maximum value in DC1 as the root cell for DPT estimation. To visualize the diffusion components in space, we added an estimated vector field to the segmented spatial graphs with arrow sizes corresponding to the magnitude of change among nearby cells, and directions corresponding to the direction with the largest change in the diffusion component. We then identified imputed genes strongly correlated with DPT (absolute Spearman correlation > 0.5) amongst either Midbrain or Hindbrain region cells. For smooth expression estimation along the DPT, we split cells into either Midbrain or Hindbrain regions and extracted fitted values from local regression (loess) for each gene with DPT ranking as the explanatory variable. To further identify genes associated with spatial variation in expression, we performed scHOT^82^ analysis using weighted mean as the underlying higher order function, with a weighting span of 0.1 on spatial coordinates and using the imputed gene expression values. We then identified the 500 top-ranked significantly spatially variable genes (ensuring also that FDR-adjusted P-value < 0.05), and clustered their smoothed expression using hierarchical clustering (Supplementary Table 7), selecting the number of clusters using dynamicTreeCut^120^. To visualize spatial expression profiles of clusters, we calculated the mean inferred gene expression value for the genes associated with each cluster.

### Joint analysis with Nowotschin et al. (2019) dataset

We downloaded the Nowotschin *et al*. 10X Genomics scRNA-seq counts and associated annotations from the corresponding Shiny web application (https://endoderm-explorer.com/)^2^. We then subset down to E8.75 cells, considering each 10X Genomics sequencing library as a batch variable. We performed highly variable gene (HVG) selection using ‘modelGeneVar’ from the scran package^108^, using the library sample as the blocking variable. We then selected the intersection of these HVGs and the genes appearing in the seqFISH dataset for further analysis. We concatenated the normalized logcounts for the Nowotschin *et al.* and seqFISH datasets and performed dimensionality reduction to 50 principal components using ‘multiBatchNorm’ as implemented in scran^108^. We then performed batch correction using the Mutual Nearest Neighbors approach, where the merge order was fixed to first integrate batches from the Nowotschin *et al.* dataset (ordered by decreasing cell number). We then identified the 10 nearest neighbors of the seqFISH cells to the Nowotschin *et al.* cells in the corrected reduced dimensional space. Using these nearest neighbors, we classified seqFISH Gut tube cells to a cell type defined by Nowotschin *et al..* A “mapping score” was computed for each cell as the proportion of the nearest neighbors in Nowotschin *et al.* data corresponding to the selected class. We performed differential gene expression analysis between the Lung 1 and Lung 2 groups using ‘findMarkers’ in scran^108^, and also performed differential gene expression analysis between the associated mesodermal cells at most three steps away from the Lung 1 or Lung 2 cells in the cell-cell neighborhood network.

### Anterior-Posterior axis cell ranking

To calculate the relative position of developing gut tube cells along the anterior-posterior axis, for each embryo we performed a virtual dissection to visually identify the dorsal and ventral regions of the gut tube. Then for each embryo and each dorsal or ventral tissue region, we fit a single principal curve model, using the R package princurve^121^, with explanatory variables corresponding to the physical coordinates. We then extracted anterior-posterior cell rankings by taking the rank of the fitted arc-length from the beginning of the curve, ensuring the curve always began at the anterior-most position.

### Joint analysis with Nowotschin et al. (2019) and Han et al. (2020) datasets

To further understand the relationship between the endodermal and mesodermal layers in the gut tube, we performed joint analysis between the Nowotschin *et al.* data (described above), as well as the E8.5 splanchnic mesoderm cells from Han *et al.*^3^. For the Han *et al*. data, we performed highly variable gene (HVG) selection using ‘modelGeneVar’ from the scran package^108^, using the library sample as the blocking variable, and then selected the genes that appeared in either the HVG list for Nowotschin *et al.* or Han *et al.*, and that were also present in the seqFISH gene library. We then concatenated the normalized logcounts of all three datasets and performed integration (dimensionality reduction, batch correction, further dimensionality reduction for visualization) and cell classification as described above. Thus, for each seqFISH cell, we obtained a classified cell class according to the labels provided by Han *et al.*, including mesodermal subtypes in the splanchnic mesoderm. To further investigate the surrounding mesodermal cells of the gut tube, we used the cell-cell neighborhood network to identify mesodermal cells at most three steps away from a gut tube cell and, for each of these cells, we identify their position as either dorsal or ventral to the gut tube, and calculated the mean position along the anterior to posterior axis.

## Data availability

The spatial transcriptomic map can be explored interactively at: https://marionilab.cruk.cam.ac.uk/SpatialMouseAtlas/ and raw image data is available on request. Processed gene expression data with segmentation information and associated metadata is also available to download and explore online at https://marionilab.cruk.cam.ac.uk/SpatialMouseAtlas/. Scripts for downstream analysis are available at https://github.com/MarioniLab/SpatialMouseAtlas2020.

## Acknowledgements

We thank our colleagues in the Wellcome Trust Mouse Gastrulation Consortium, as well as colleagues in the University of Cambridge Stem Cell Institute, Cancer Research UK Cambridge Institute, Babraham Institute, Gurdon Institute, California Institute of Technology, Sloan Kettering, and the Francis Crick Institute for their support and intellectual engagement. We thank Julian Thomassie, Elsy Buitrago-Delgado, Jina Yun, Michael Lawson, Christopher Cronin, Chris Karp and other members of the Cai Lab for experimental support and constructive input. We thank Sonja Nowotschin and Esther Wershof for discussions concerning the gut tube analysis. We thank Veronique Juvin for providing scientific illustration. We thank Daniel Keitley and Michael Morgan and other members of the Marioni Lab for discussions concerning the analysis. We thank members of the Nichols and Reik lab for their discussions concerning the project.

## Author Contributions

T.L., J.A.G. and C.G., performed the probe library gene selection, with input from W.R., J.N., B.G. and J.C.M.. T.L., E.S.B. and J.N. performed embryo collection. T.L., C.-H.L.E and N.K. performed the seqFISH method and segmentation optimization for mouse embryonic tissue sections. T.L. and N.K. generated the spatial dataset, with supervision from L.C.. N.P. performed image processing including registration, cell segmentation and mRNA spot decoding, with input from L.C.. T.L. and A.M. performed optical threshold selection for non-barcoded smFISH images. S.G. performed pre-processing, low-level analyses, batch correction, clustering, integration with other datasets, global visualization, and designed the associated website, with input from A.M., R.A.. A.M. performed imputation analysis. S.G. and A.M. performed analysis surrounding midbrain hindbrain boundary formation. E.S.B. performed HCR imaging experiments, with supervision from A.-K.H.. R.T., C.G., S.S., J.B., B.D.S., A.-K.H., B.G., W.R. and J.N. provided discussion and interpretation of the data and analysis. B.G., W.R., J.N., L.C., and J.C.M. supervised the study. T.L., S.G. and A.M. generated figures. T.L., S.G., A.M. and J.C.M. wrote the manuscript. L.C. and J.C.M. oversaw the entirety of the project. All authors read and approved the final manuscript.

