## Supplementary Material for "Highly multiplexed spatially resolved gene expression profiling of mouse organogenesis"

### SUPPLEMENTARY FIGURES

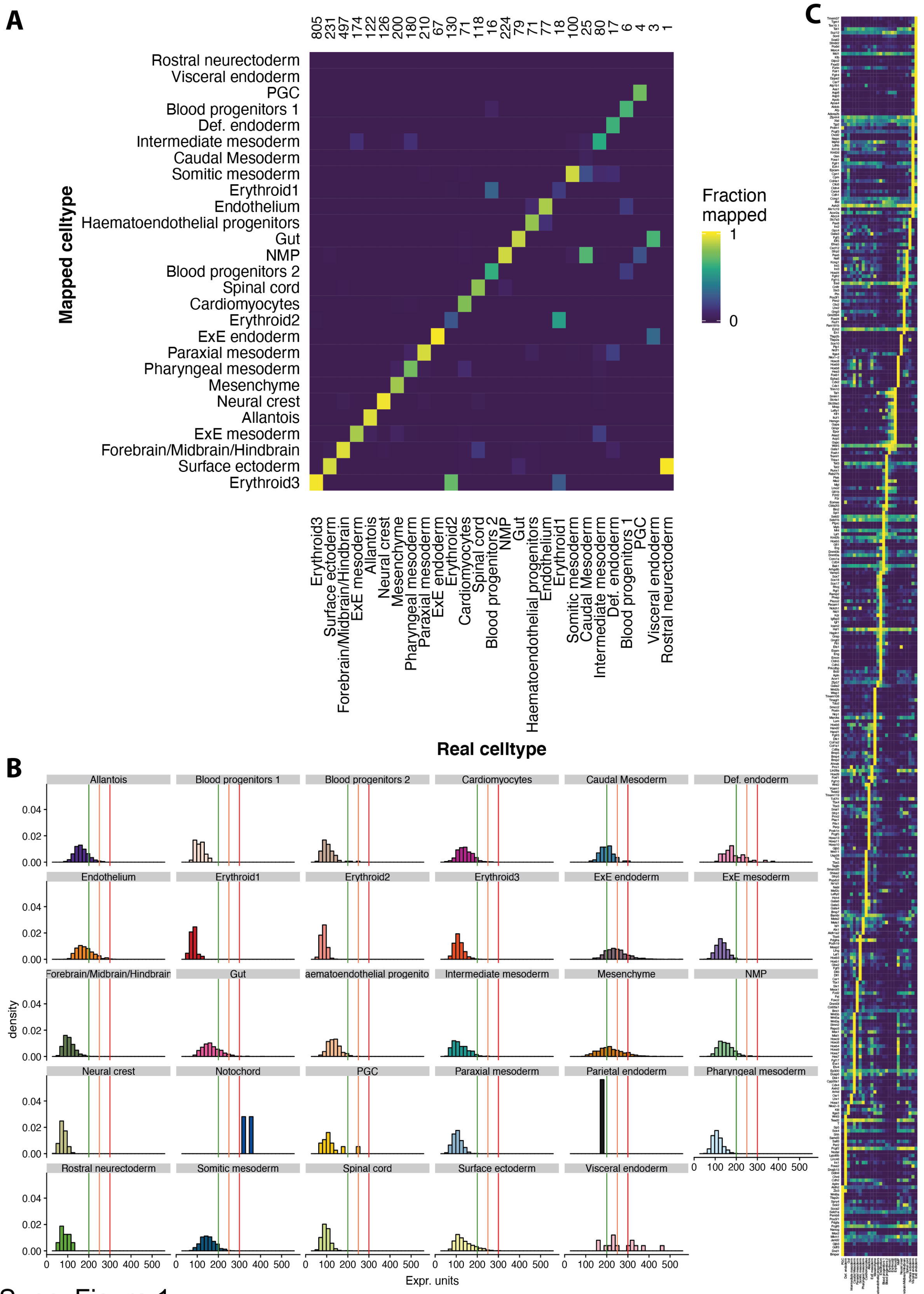


Supp. Figure 1

**Supplementary Figure 1: seqFISH probe library design**

(A) Predicting Gastrulation atlas cell types using the seqFISH probe library. The x-axis is the true cell type of each cell, and the y-axis the mapped cell type. Shading indicates the fraction of cells of each true cell type mapped to each possible cell type. Numbers for each column correspond to the number of cells in each true cell type.

(B) Histogram, showing the seqFISH library feasibility. Histograms of expression units of the seqFISH probe library genes for each cell type in the E8.5 Gastrulation atlas. Green, orange, and red lines correspond to 200, 250, and 300 normalized expression units respectively, reflecting the guided expression to avoid oversaturation.

(C) Heatmap showing the mean expression of all selected seqFISH library genes (rows) for each cell type (columns) in the E8.5 Gastrulation atlas.

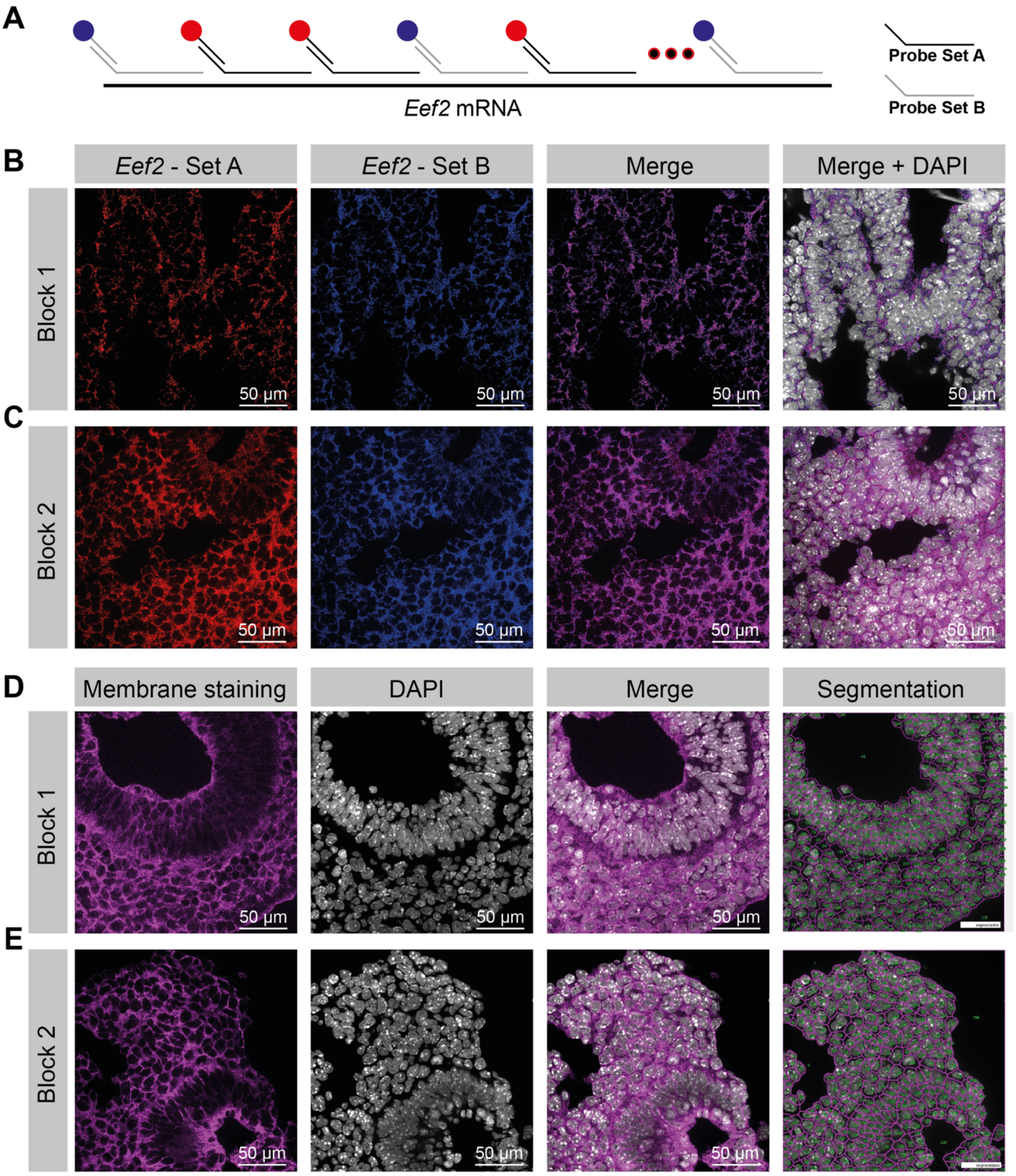


Supp. Figure 2

**Supplementary Figure 2: Validation of RNA quality and cell segmentation**

(A) Schematic overview of the hybridization of two interspersed *Eef2* probe sets to test for RNA integrity.

(B) Image showing the expression of *Eef2* probe set A (Alexa Fluor 647 - red) and *Eef2* probe set B (Cy3B - blue) for experimental block 1. Color merge of these two images indicates a high degree of overlap between red and blue probes. Merge and DAPI (grey) show overlap of *Eef2* signal surrounding regions where cell nuclei are present.

(C) Expression profile of *Eef2* probe set A and B, as described in (B) for experimental block 2.

(D) Image of cell membrane labeling (purple) using a combination of E-cadherin, N-cadherin, Pan-cadherin and β-catenin primary antibody staining, following an optimized cell segmentation protocol (Methods) and nuclear staining using DAPI (grey) for the first tissue section, containing embryo 1 and 2. Signal membrane labeling was used for cell segmentation using Ilastik[36].

(E) Cell membrane labeling (purple) and cell segmentation, as described in (D) for experimental block 2.

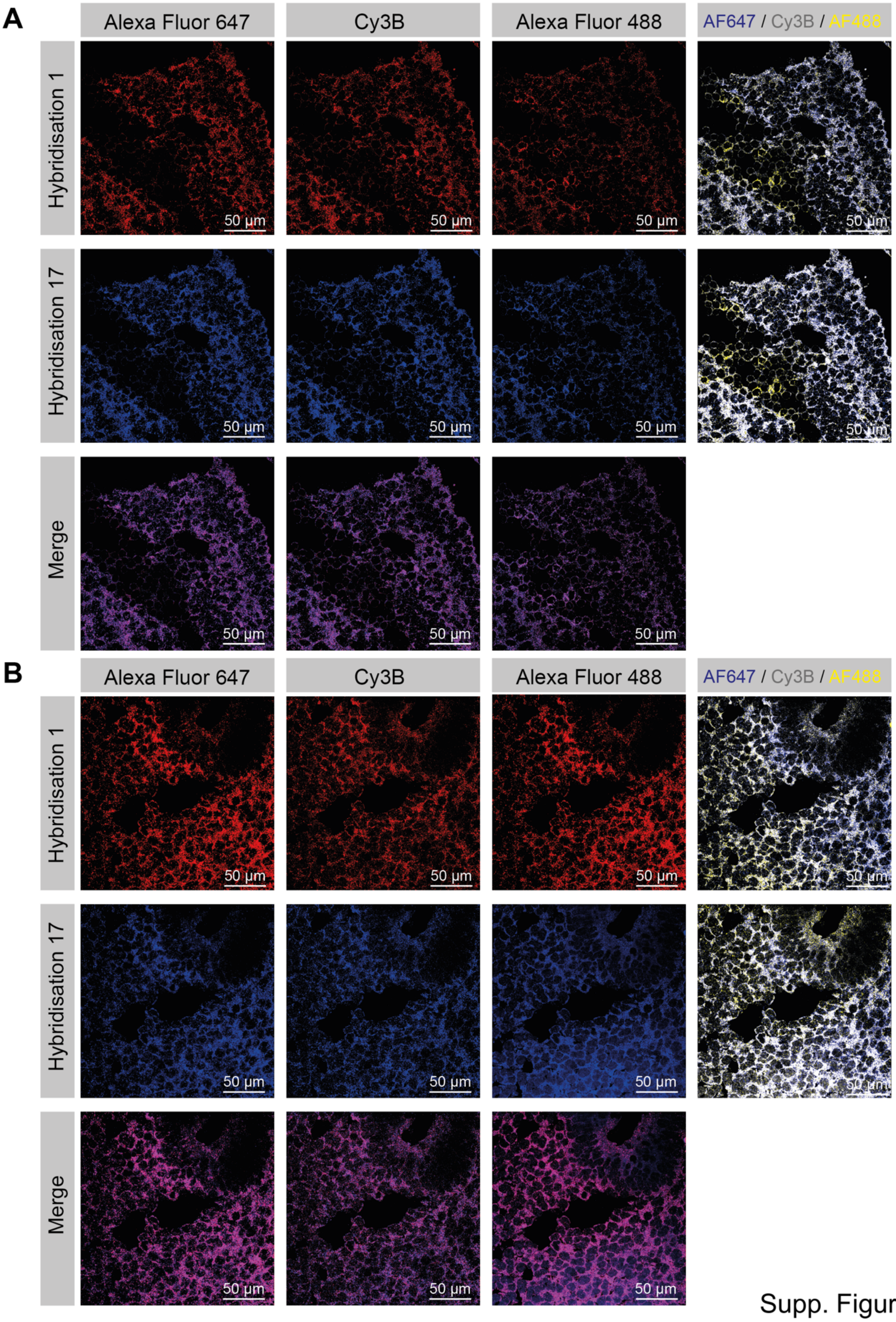


Supp. Figure 3

**Supplementary Figure 3: Comparison of the hybridization round 1 and 17, an additional repeat of hybridization round 1, for quality control.**

(A) Visualization of the experimental block 1, containing embryos 1 and 2, and the experimental block 2 (B), containing embryo 3. mRNA spots for the Alexa fluor 647, Cy3B and Alexa fluor 488 channels are shown separately for hybridization round 1 (red) and 17 (blue). Strong overlap (purple) of the mRNA spots between hybridization rounds 1 and 17 suggests high RNA quality after the seqFISH imaging. A composite of all three channels is visualized for both hybridization rounds separately.

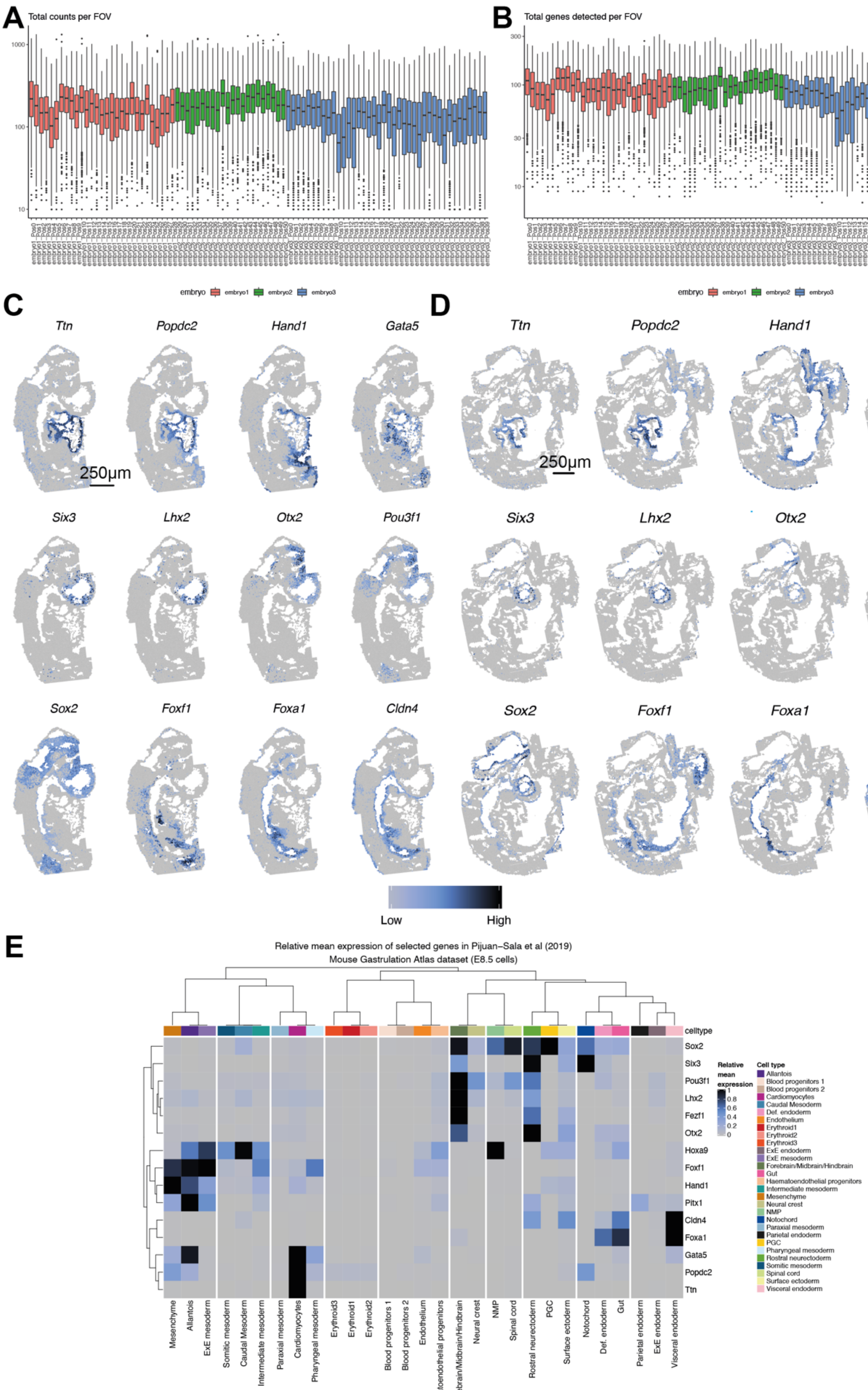


Supp. Figure 4

**Supplementary Figure 4: Quality Control of seqFISH data.**

(A) Boxplot of total mRNA molecules detected for each field of view (log10 scale), colored by embryo.

(B) Total number of genes detected for each field of view (log10 scale), colored by embryo.

(C) Spatial expression of selected genes shown in Figure 1 for embryo 2. Scale bar 250 µm.

(D) as in C for embryo 3. Scale bar 250 µm.

(E) Heatmap of relative mean expression of cell types corresponding to E8.5 Gastrulation atlas data, cell type dendrogram corresponds to clustering using all data.

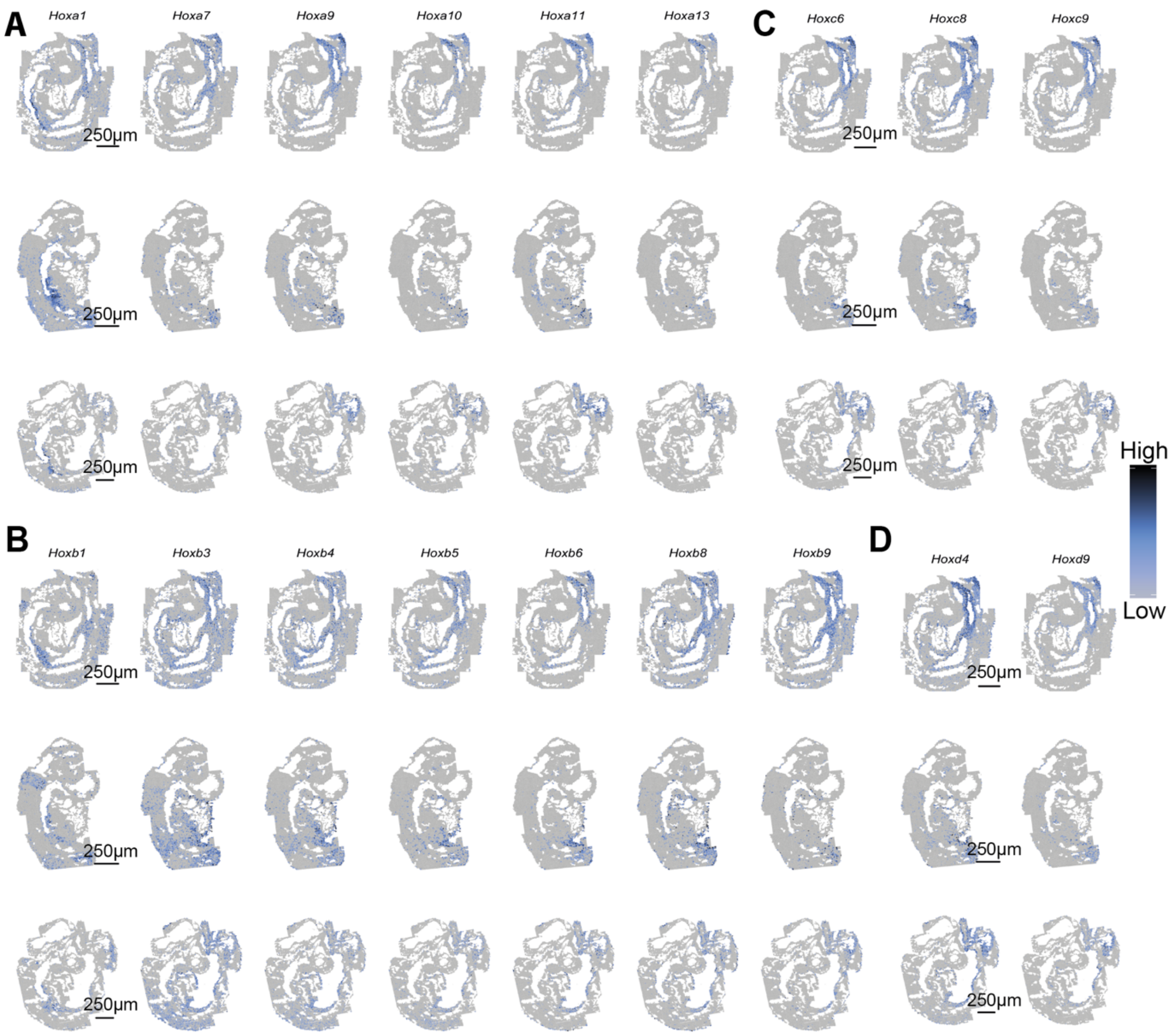


Supp. Figure 5

**Supplementary Figure 5: Spatial Hox expression profiles to assess data quality.**

(A) Spatial expression of HoxA family genes, ordered numerically, with each embryo per row. Scale bar 250 μm.

(B) as in A, with HoxB subfamily.

(C) as in A with HoxC subfamily.

(D) as in A with HoxD subfamily.

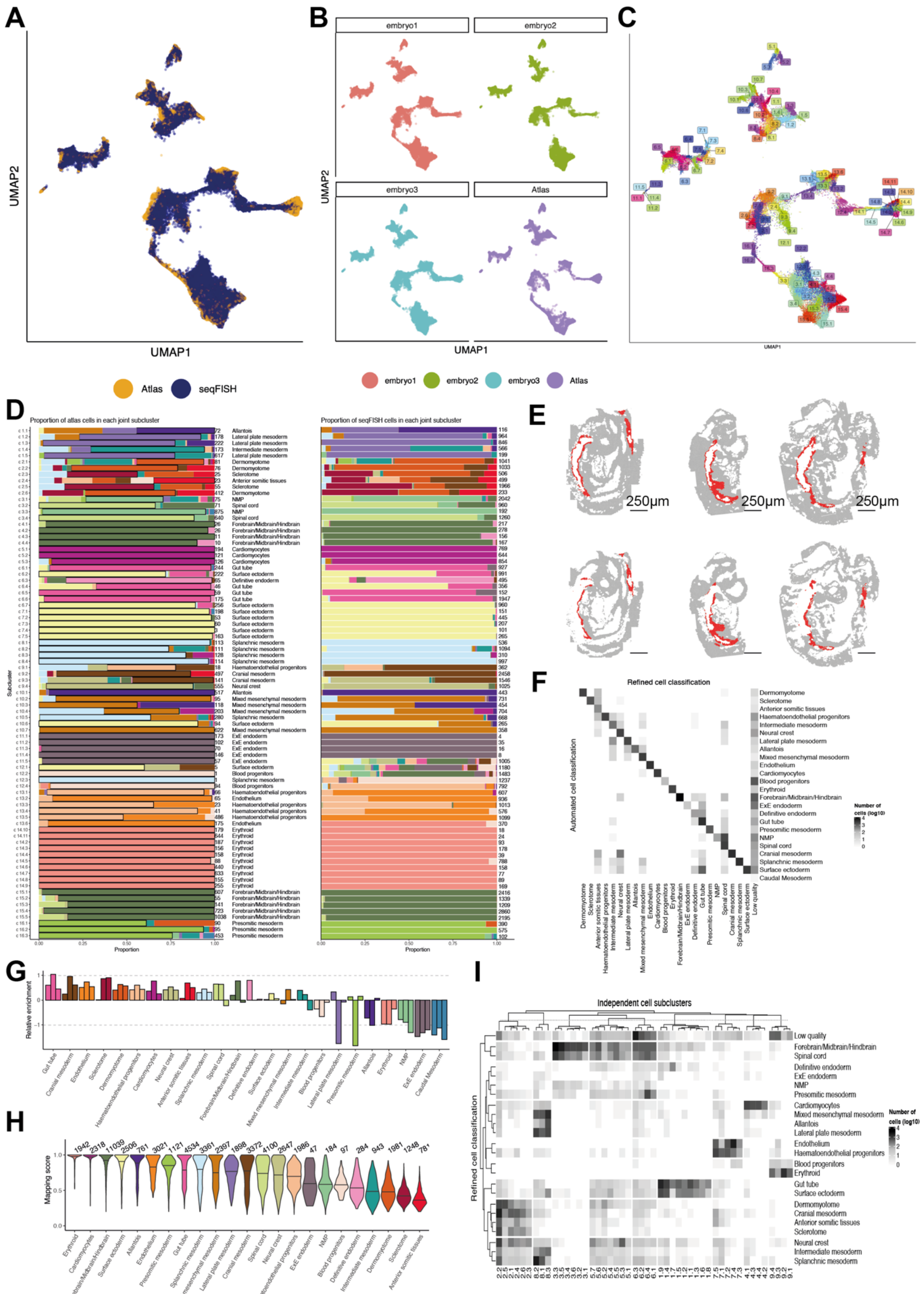


Supp. Figure 6

**Supplementary Figure 6: Optimizing cell type annotation**

(A) Joint UMAP of Gastrulation atlas and seqFISH expression data, with cells colored by data modality.

(B) Joint UMAP of Gastrulation atlas and seqFISH expression data, with panels corresponding to each embryo and the Gastrulation atlas dataset.

(C) Joint UMAP of Gastrulation atlas and seqFISH expression data, colored by joint subclustering with labels corresponding to centroid in UMAP coordinates.

(D) Barplots of the proportion of cell types from the Gastrulation atlas cells present in each subcluster (left), and automated cell type classification for seqFISH data (right). Numbers beside each bar correspond to the number of cells, and labels beside the left barplot correspond to the majority cell type of the Gastrulation atlas cells for each joint subcluster.

(E) Spatial map of virtual dissection of cells to be classed as developing gut tube, for each embryo (columns) and z-slice (rows). Scale bar 250 μm.

(F) Heatmap of contingency table of automated cell type label for seqFISH cells (rows) and refined cell type classification (columns).

(G) Barplot of relative enrichment in abundance of seqFISH cells compared to Gastrulation atlas cells, each bar corresponds to embryo 1, 2, and 3, from left to right.

(H) Violin plots of automated cell type mapping score for each seqFISH cell, with bar corresponding to median. Numbers above correspond to the number of cells classed into each cell type.

(I) Heatmap of contingency table of cell type label for seqFISH cells (rows) and independent unsupervised cell subclusters (columns).

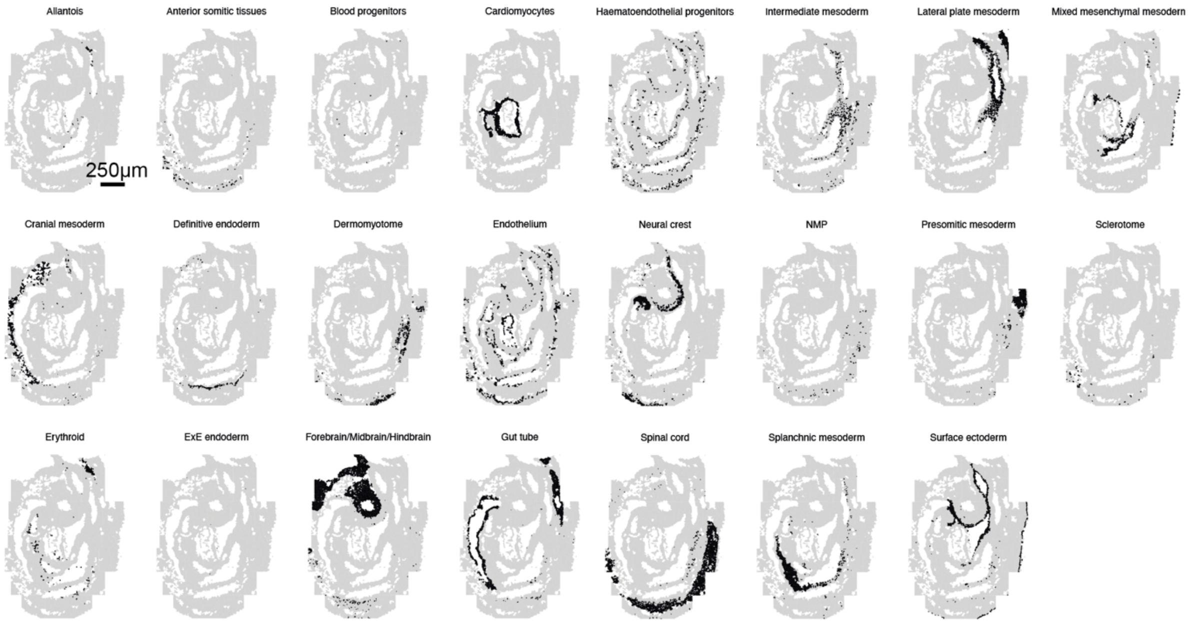


Supp. Figure 7

**Supplementary Figure 7: Cell type annotation for Embryo 1**
Spatial plots of embryo 1 where, for each panel, the selected cell type is shown in black.

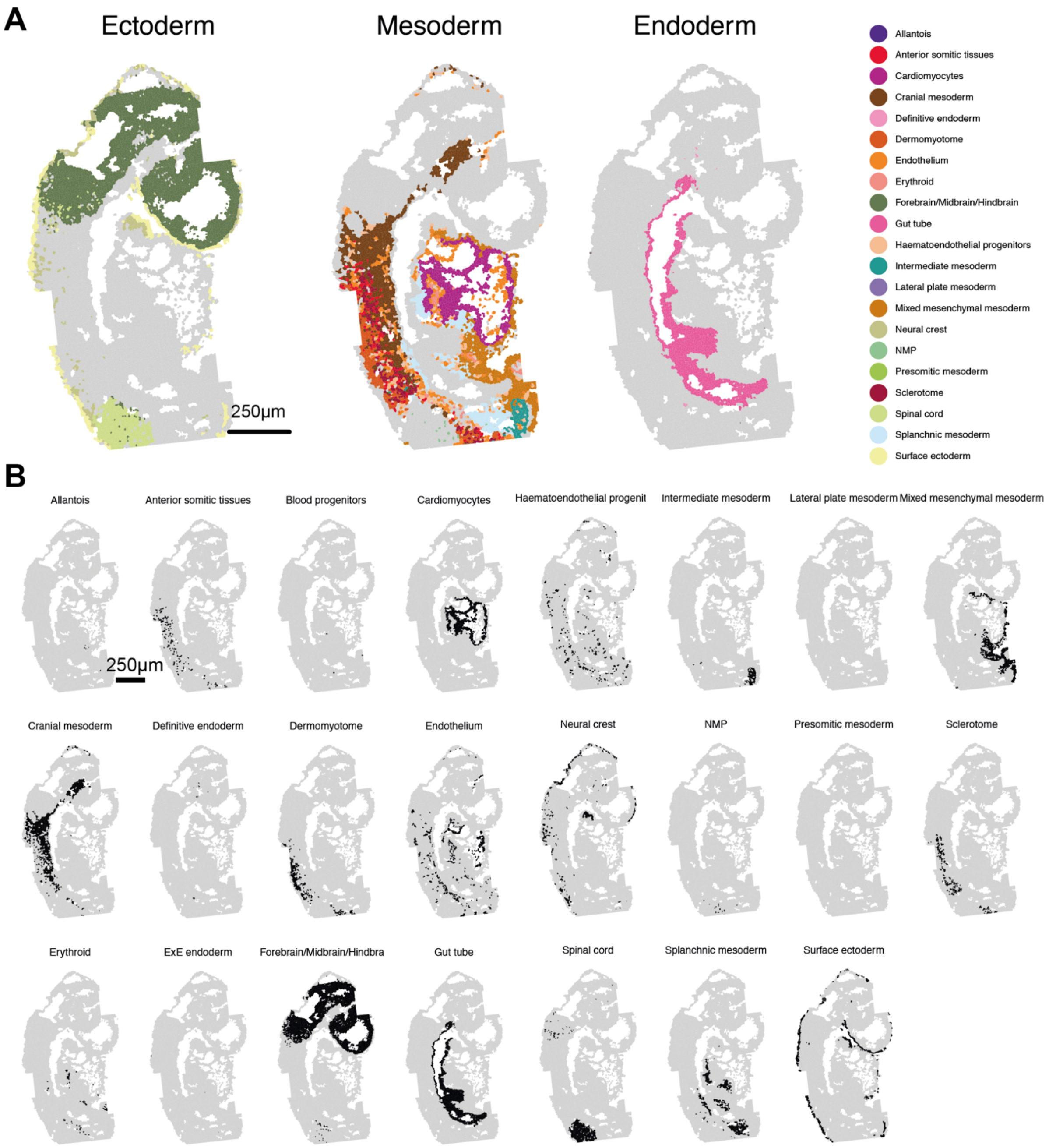


Supp. Figure 8

**Supplementary Figure 8: Cell type annotation for Embryo 2**

(A) Cell type maps separated by the three germ layers (ectoderm, mesoderm, endoderm) for embryo 2. Scale bars 250 µm.

(B) Spatial plots of embryo 2 where, for each panel, the selected cell type is shown in black.

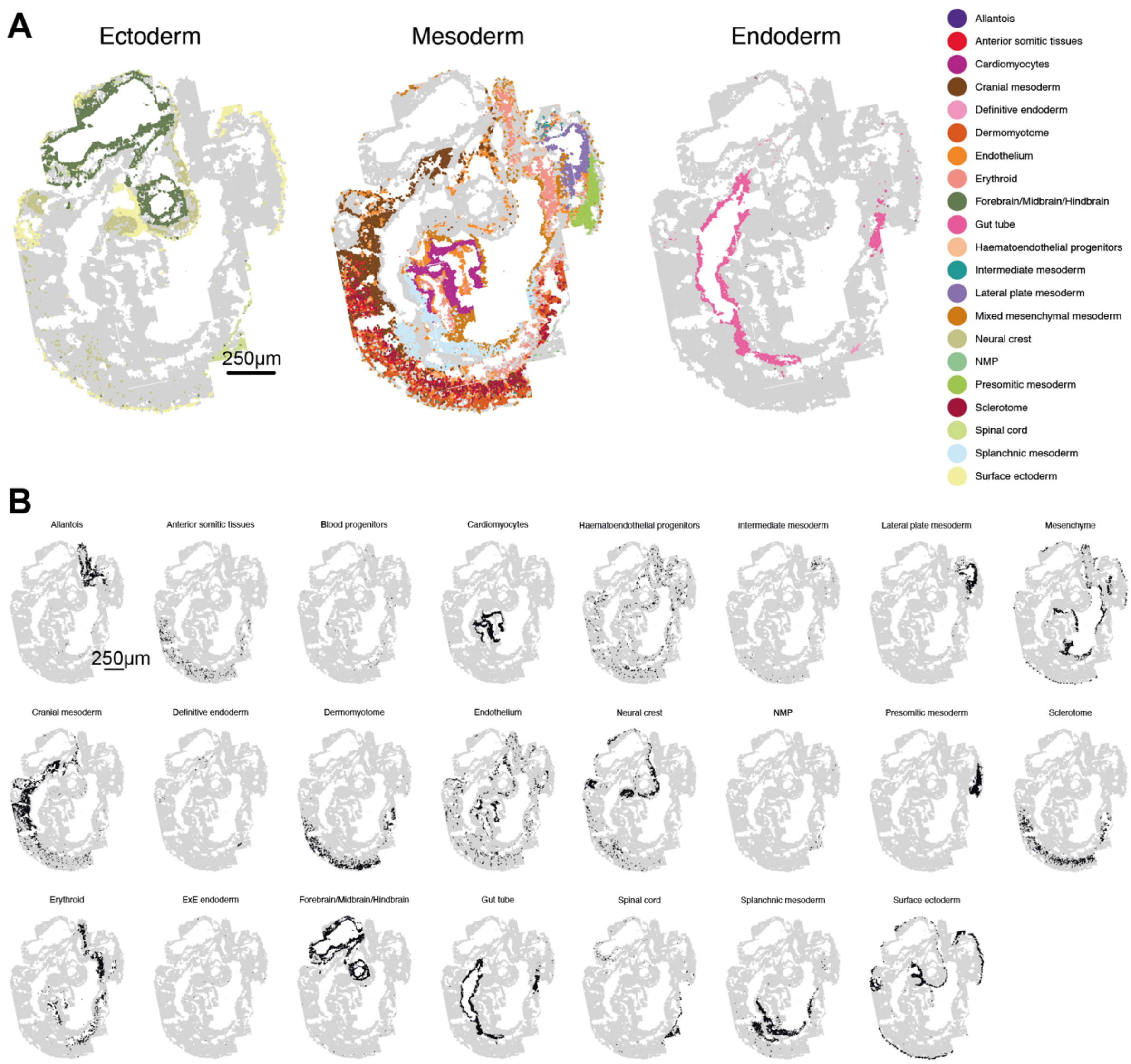


Supp. Figure 9

**Supplementary Figure 9: Cell type annotation for Embryo 3**

(A) Cell type maps separated by the three germ layers (ectoderm, mesoderm, endoderm) for embryo 3. Scale bars 250 µm.

(B) Spatial plots of embryo 3 where, for each panel, the selected cell type is shown in black.

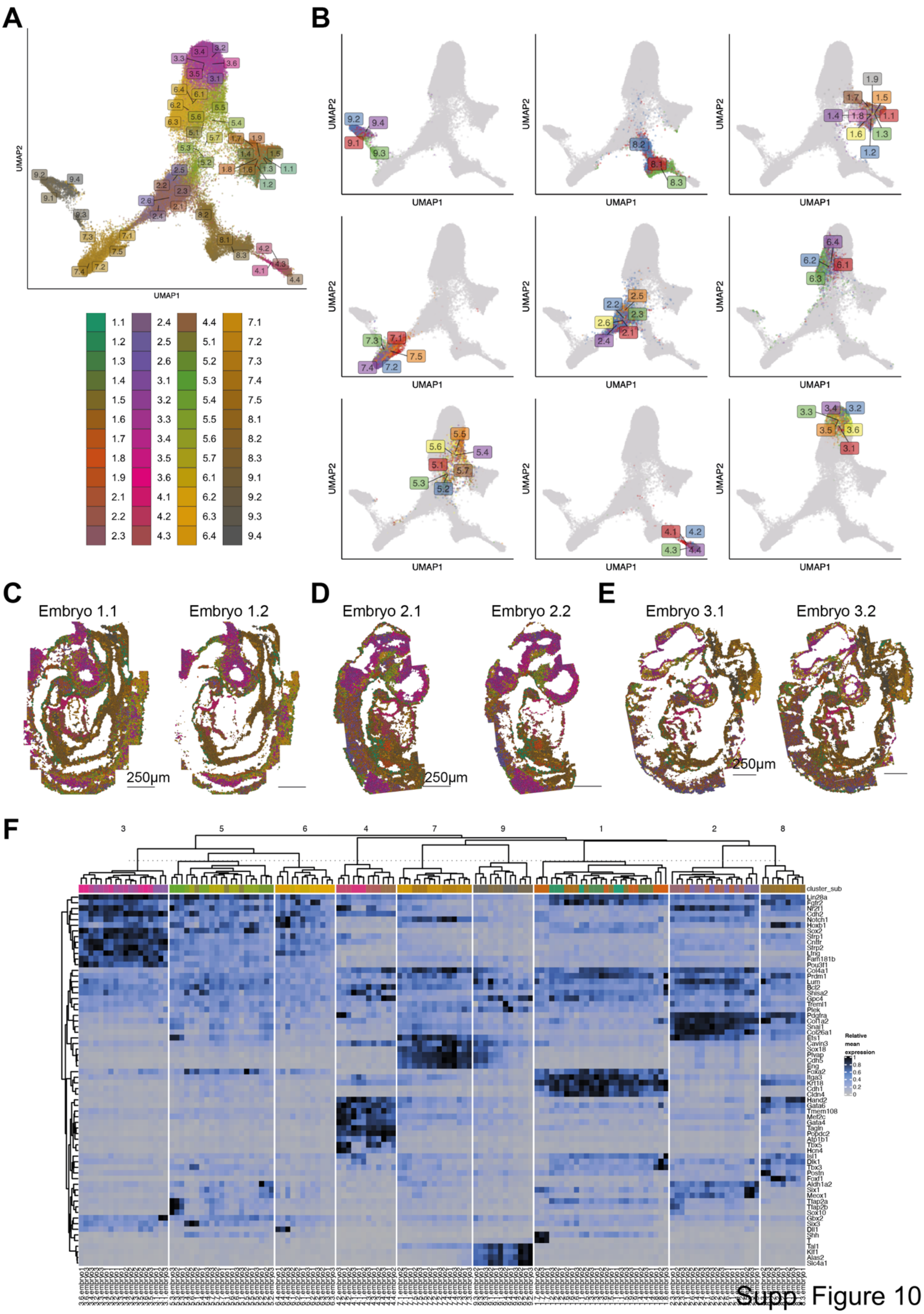


Supp. Figure 10

**Supplementary Figure 10: Unsupervised clustering of seqFISH data**

(A) UMAP of seqFISH expression data, with cells colored by unsupervised subclusters, with labels corresponding to centroid in UMAP coordinates.

(B) Multiple panels displaying UMAP of seqFISH expression data, with cells for each separate cluster colored by the associated subcluster, with labels corresponding to centroid in UMAP coordinates.

(C) Spatial map of embryo 1 cells colored by unsupervised subclusters (colors matching panel A) for each z-slice. Scale bar 250 μm.

(D) as in C with embryo 2.

(E) as in C with embryo 3.

(F) Heatmap of relative mean expression of seqFISH cells grouped by embryo and unsupervised subcluster (columns) for genes selected as appearing in the top three significant marker genes (rows) for any of the subclusters. Colors along the top correspond to unsupervised subclusters with legend matching panel A.

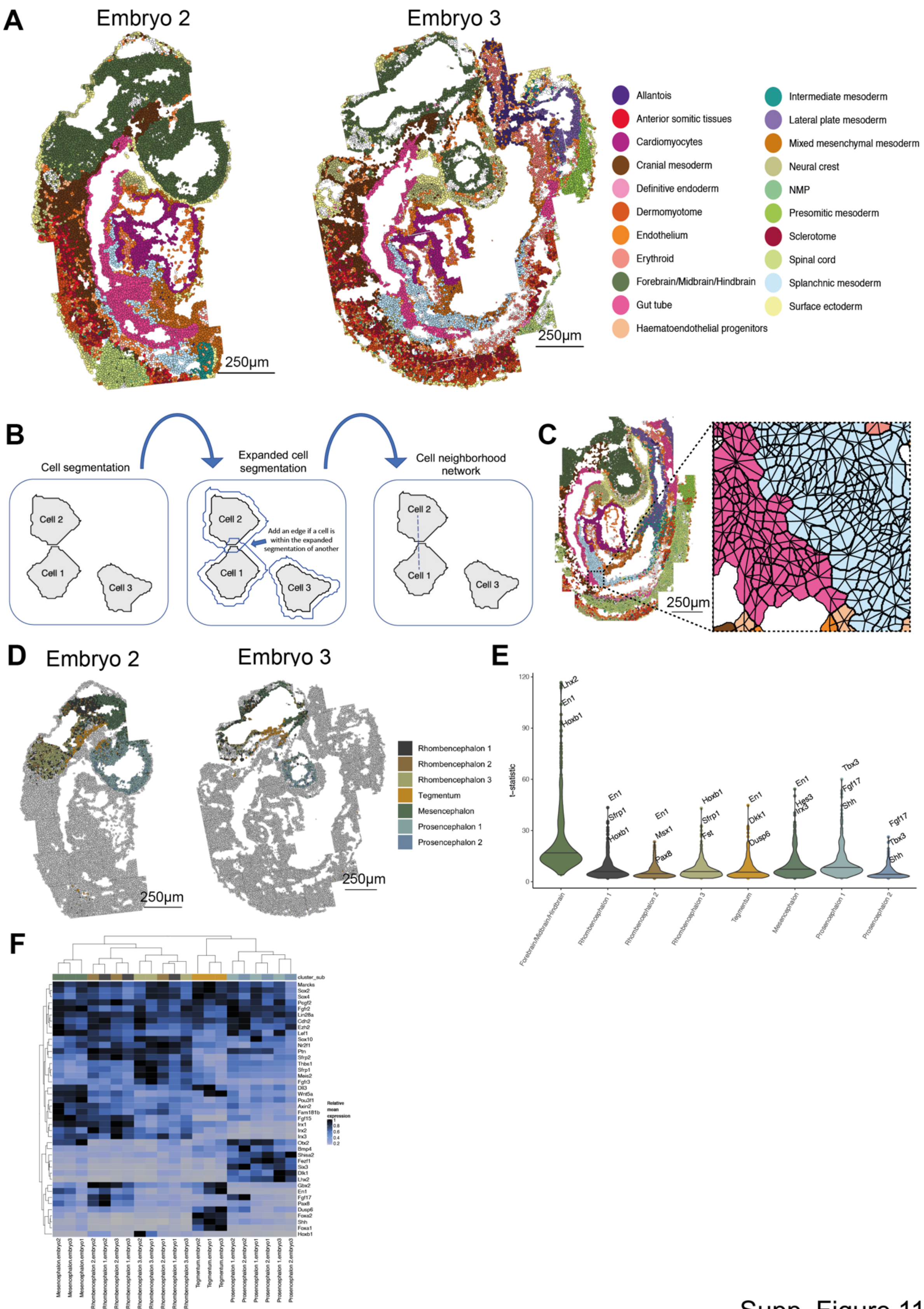


Supp. Figure 11

**Supplementary Figure 11: Cell annotation and constructing the cell-contact map**

(A) Spatial map of embryos 2 and 3, colored by refined cell type. Scale bar 250 µm.

(B) Schematic of construction of cell neighborhood network, where cell segmentation polygons are expanded and a network edge drawn if another cell is within the expanded polygon region.

(C) Visualization of cell neighborhood network using spatial map of embryo 1 with zoom in to reveal cell neighborhood network edges among cells. Scale bar 250 µm.

(D) Spatial maps of embryos 2 and 3, with cells colored by brain subtypes, and other cells in grey. Scale bar 250 µm.

(E) Violin plot showing t-statistic corresponding to spatial heterogeneity test for each gene within brain subtype. The top three genes are labeled for each violin, and the bar corresponds to the median.

(F) Heatmap of relative mean expression of each embryo and brain subcluster for significant (FDR-adjusted P-value $< 0.05$, absolute LFC $> 0.2$) marker genes.

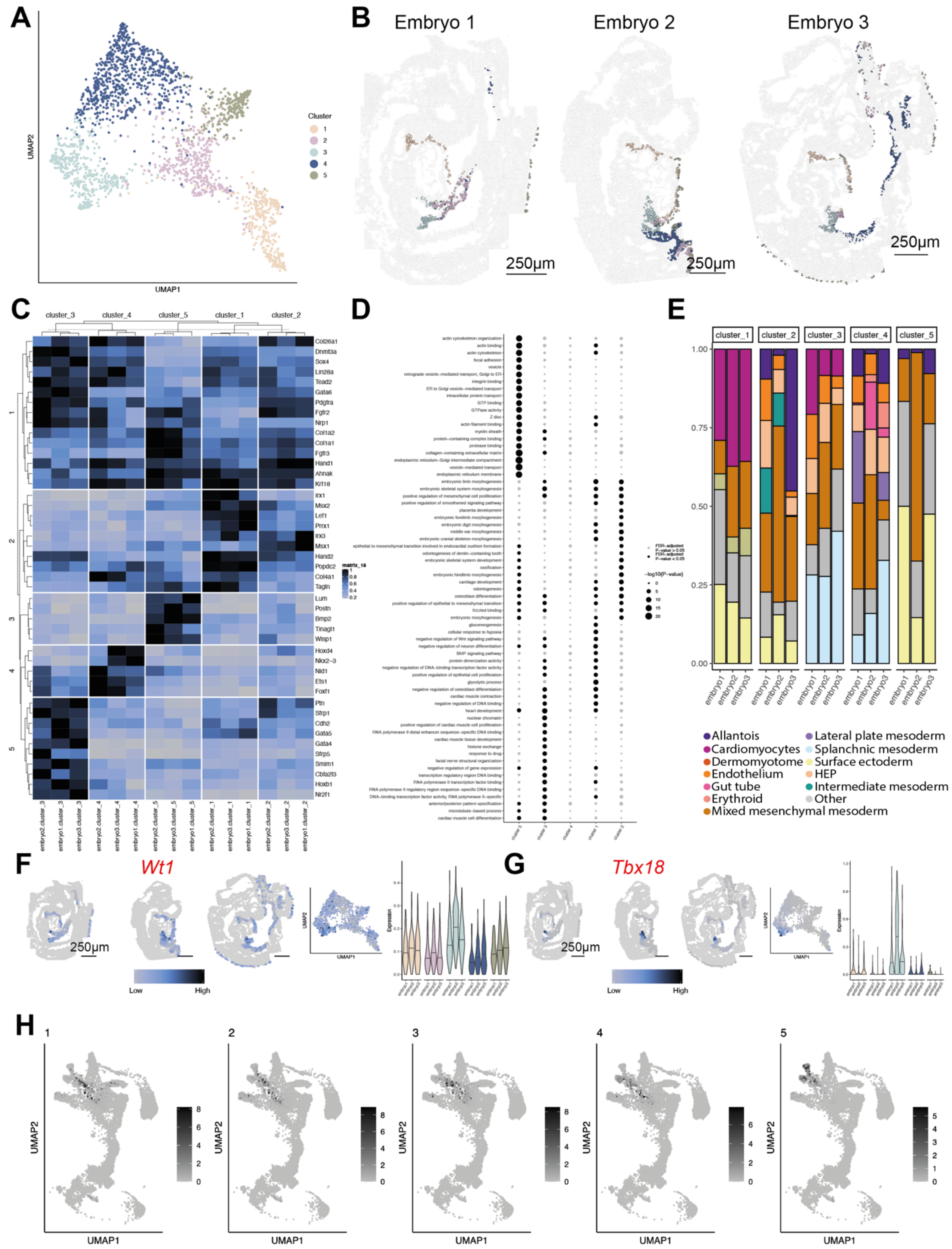


Supp. Figure 12

**Supplementary Figure 12: Characterization of mixed mesenchymal mesoderm cluster**

(A) UMAP embedding of mixed mesenchymal mesoderm seqFISH cells, colored by unsupervised clusters.

(B) Spatial plots with cells colored by mixed mesenchymal mesoderm unsupervised clusters.

(C) Heatmap of mean expression of each embryo and mixed mesenchymal mesoderm cluster for significant (FDR-adjusted P-value < 0.05, absolute LFC > 0.2) marker genes.

(D) Dotplot of significantly enriched gene ontology terms for each mixed mesenchymal mesoderm cluster.

(E) Proportional bar plot showing the corresponding cell types for spatial neighbors of each embryo and mixed mesenchymal mesoderm cluster, with cell types with a small percentage grouped into Other cell types.

(F) Spatial plots of inferred *Wt1* expression among mixed mesenchymal mesoderm clusters, UMAP embedding of cells colored by Wt1 expression, and violin plot of Wt1 expression for each embryo and mixed mesenchymal mesoderm cluster.

(G) As for (F) for inferred expression of *Tbx18*.

(H) Scatterplot of UMAP embedding of E8.5 Gastrulation atlas cells, colored by proportion of selection within nearest neighbor set for each mixed mesenchymal mesoderm cluster. Abbreviation used: HEP = haematoendothelial progenitors.

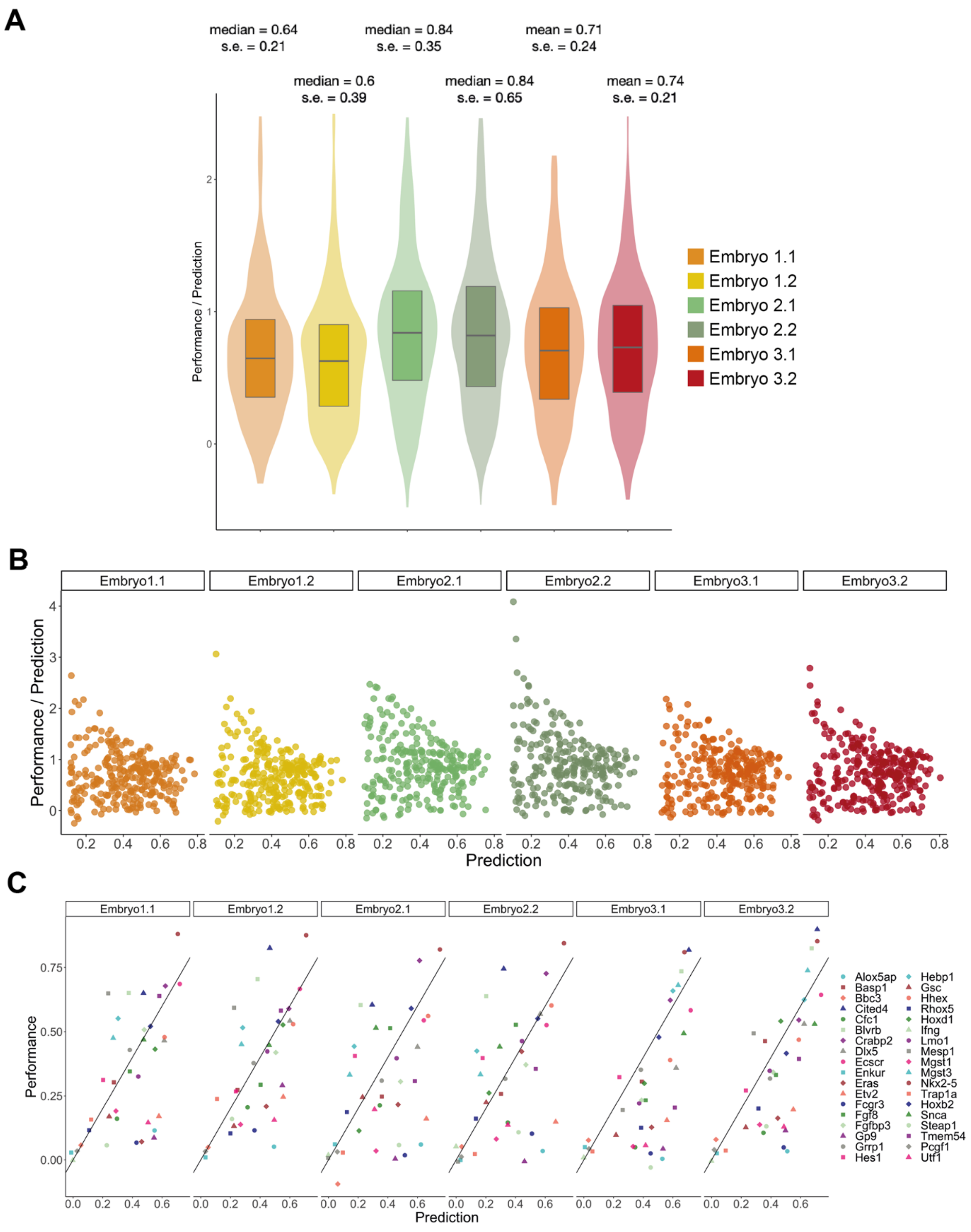


Supp. Figure 13

**Supplementary Figure 13: Imputation strategy**

(A) Normalized performance as a validation of imputation. Violin plots show distributions (across measured genes) of normalized performance for each embryo and z-slice. Median and standard error appear above each violin.

(B) Scatterplots of prediction scores (x-axis) and normalized performance scores (y-axis). Genes with prediction score lower than 0.1 show stochastic deviations in normalized performance and were filtered. Black line corresponds to running median (window size = 50) and indicates no relationship between prediction and normalized performance.

(C) Scatterplots of performance and prediction scores for genes probed by smFISH, with each panel corresponding to one embryo and z-slice, and points corresponding to genes. Genes exhibiting strong field of view effect (FOV: 39, 40, 44) were discarded from quantification of performance and prediction scores.

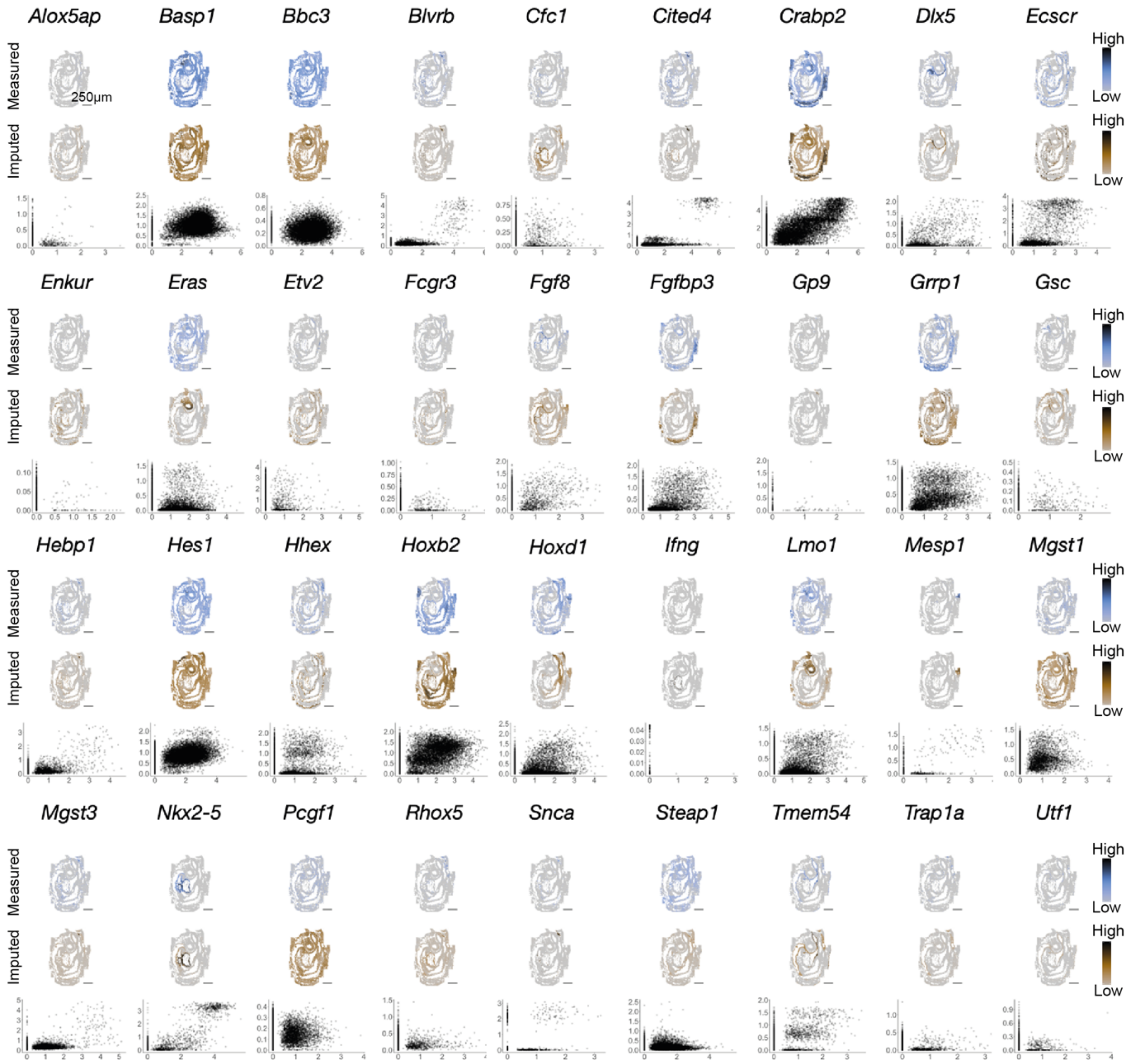


Supp. Figure 14

**Supplementary Figure 14: Comparison between imputed expression counts and measured expression counts in embryo 1.1 for 36 genes measured with smFISH.**

Each sub-panel corresponds to a single gene (denoted at the top). Upper sub-panels correspond to spatial distribution of measured logcounts (color gradient is specific to each gene). The Middle sub-panels show spatial expression maps of imputed logcounts (color gradient is specific to each gene). In the lower sub-panels, scatterplots show the measured logcounts (x-axis) and imputed logcounts (y-axis), where each point corresponds to a cell. Scale bars 250um.

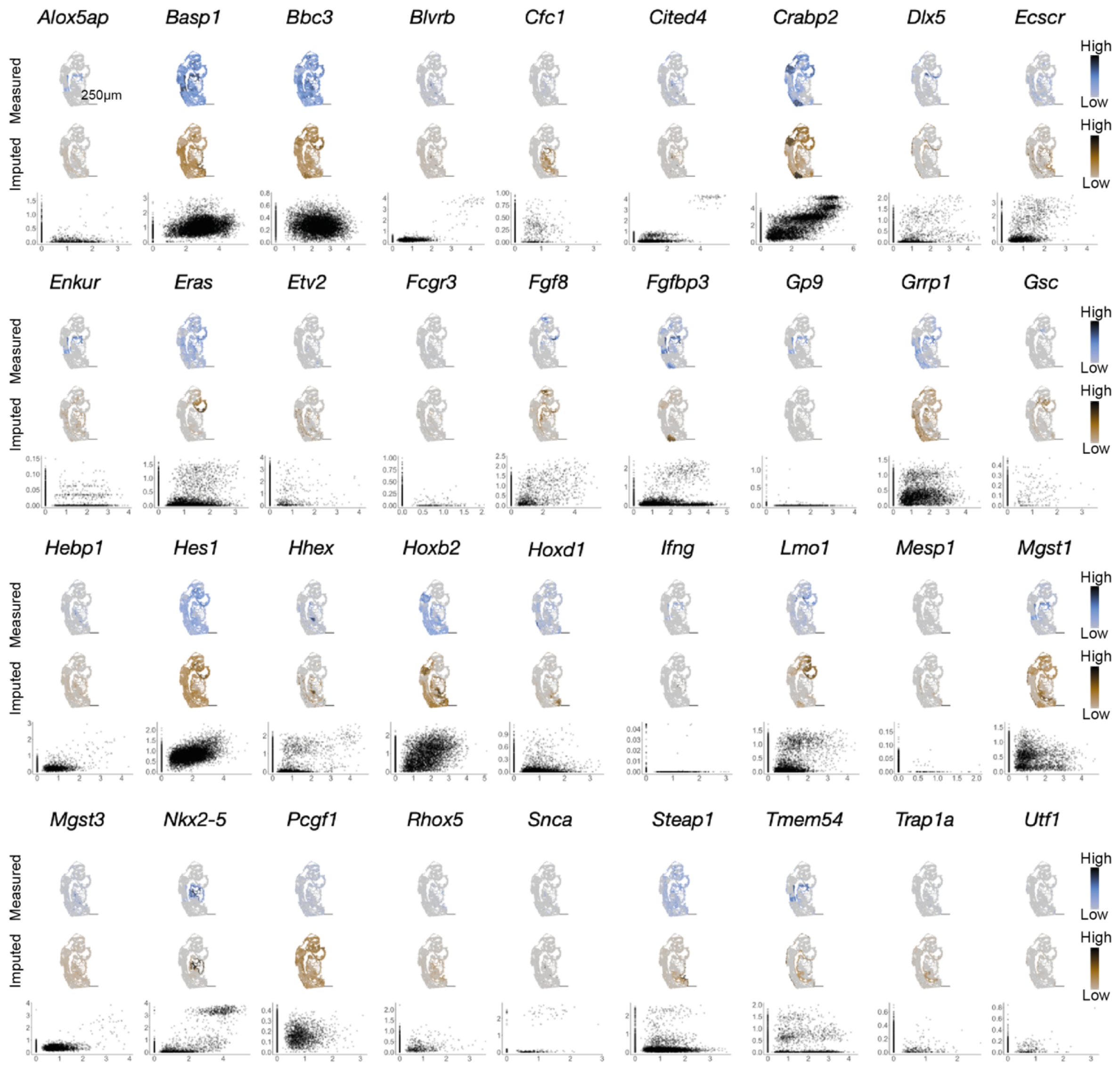


Supp. Figure 15

**Supplementary Figure 15: Comparison between imputed expression counts and measured expression counts in embryo 2.1 for 36 genes measured with smFISH.**

Each sub-panel corresponds to a single gene (denoted at the top). Upper sub-panels correspond to spatial distribution of measured logcounts (color gradient is specific to each gene). The Middle sub-panels show spatial expression maps of imputed logcounts (color gradient is specific to each gene). In the lower sub-panels, scatterplots show the measured logcounts (x-axis) and imputed logcounts (y-axis), where each point corresponds to a cell. Scale bars 250um.

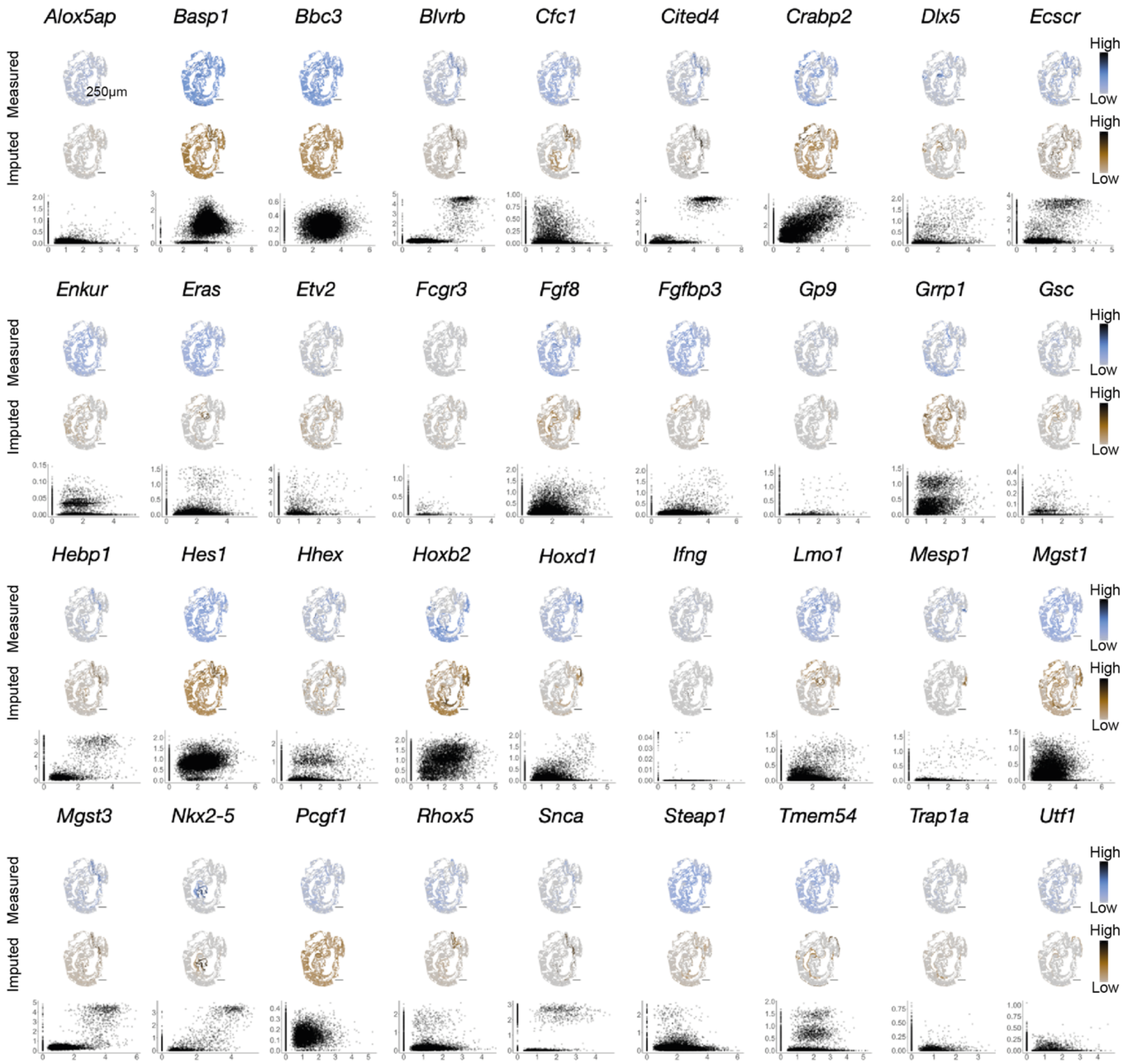


Supp. Figure 16

**Supplementary Figure 16: Comparison between imputed expression counts and measured expression counts in embryo 3.1 for 36 genes measured with non-barcoded smFISH.**

Each sub-panel corresponds to a single gene (denoted at the top). Upper sub-panels correspond to spatial distribution of measured logcounts (color gradient is specific to each gene). The Middle sub-panels show spatial expression maps of imputed logcounts (color gradient is specific to each gene). In the lower sub-panels, scatterplots show the measured logcounts (x-axis) and imputed logcounts (y-axis), where each point corresponds to a cell. Scale bars 250um.

**A**

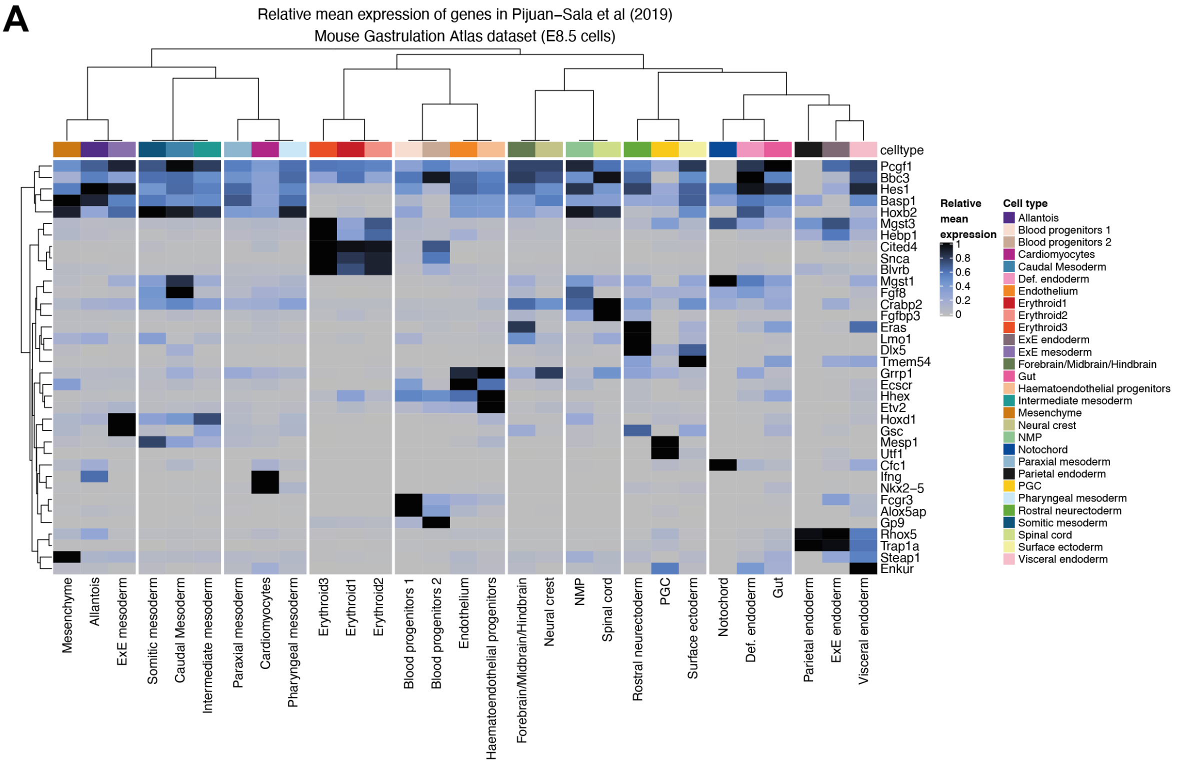


Supp. Figure 17

**Supplementary Figure 17: Expression of smFISH genes in the Gastrulation atlas**

Heatmap of relative mean expression of cell types for genes measured with non-barcoded sequential smFISH using the E8.5 Gastrulation atlas data. The cell type dendrogram was generated by clustering all data.

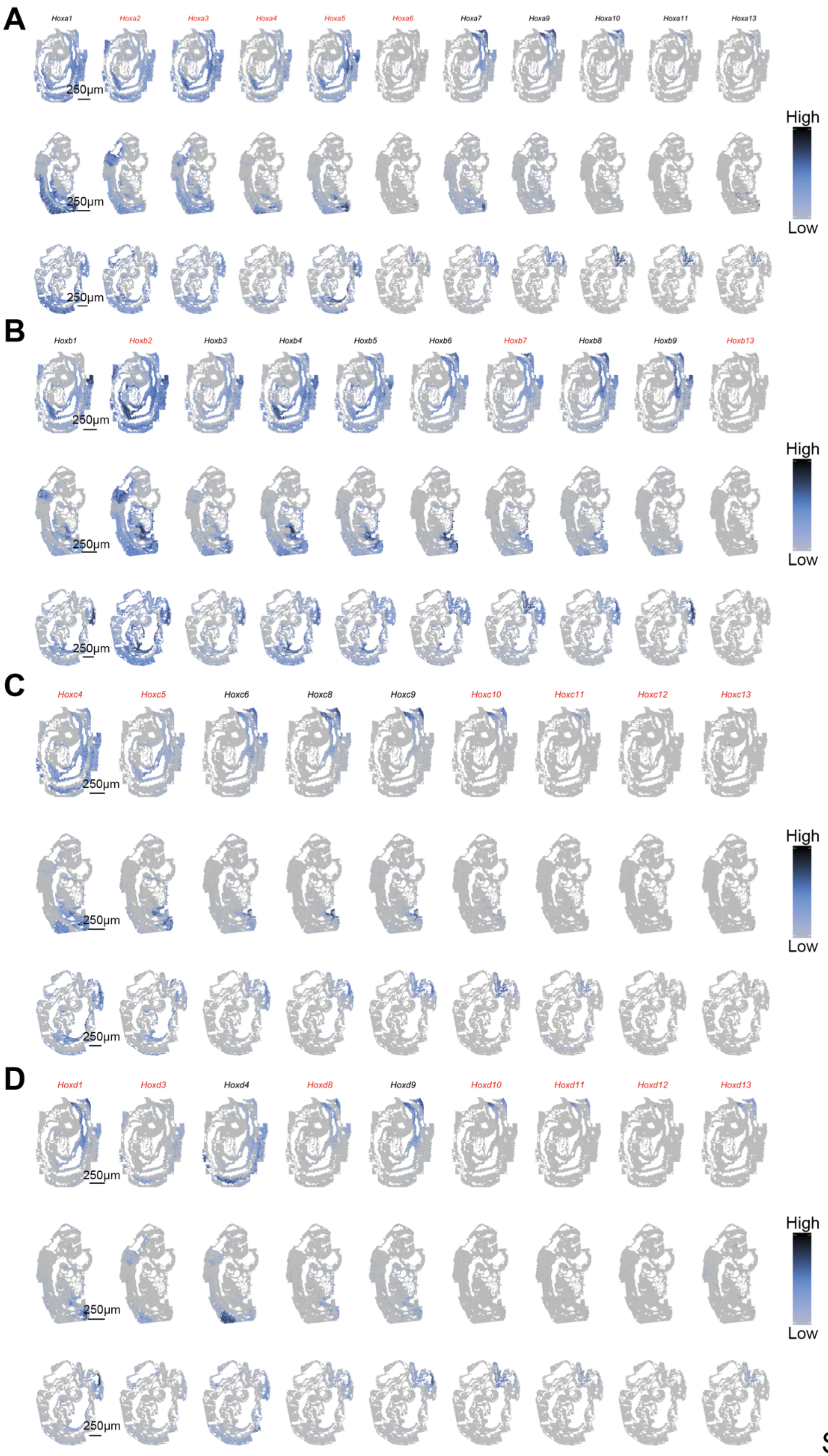


Supp. Figure 18

**Supplementary Figure 18: Spatial expression of imputed Hox gene family.**

(A) Spatial expression of all imputed HoxA family genes, ordered numerically, with each embryo per row. Gene name in red indicates whether the gene is absent from the seqFISH gene library. Scale bar 250 μm.

(B) as in A, with HoxB subfamily.

(C) as in A with HoxC subfamily.

(D) as in A with HoxD subfamily.

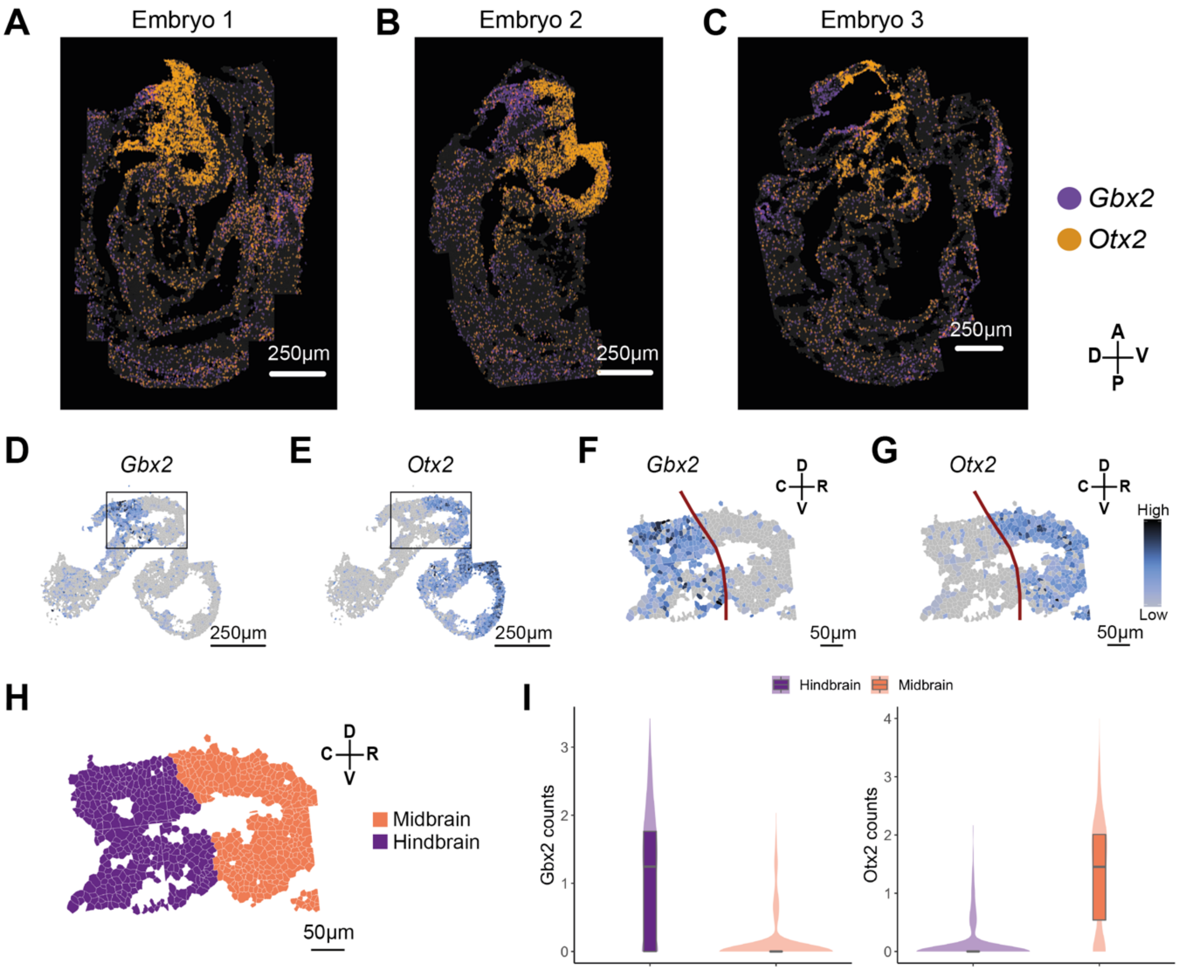


Supp. Figure 19

**Supplementary Figure 19: Virtual dissection of Midbrain-Hindbrain Boundary.**

(A) ‘Digital *in situ*’ showing detected mRNA molecules for *Gbx2* (purple) and *Otx2* (orange) across embryo 1. Scale bar 250 μm.

(B) as in A for embryo 2.

(C) as in A for embryo 3.

(D) Spatial expression of *Gbx2* in the brain. Black rectangle corresponds to the virtually dissected region in which we predict the Midbrain-Hindbrain boundary (MHB) forms. Scale bar 250 μm.

(E) as in D for the gene *Otx2*. (F) Spatial expression of *Gbx2* in the boxed region with corresponding virtual dissection (red line). Scale bar 250 μm.

(G) as in F for gene *Otx2*.

(H) Spatial distribution in the boxed region where cells are colored based on whether they are assigned a Midbrain (orange) or Hindbrain (purple) identity.

(I) Quantitative distribution of *Otx2* and *Gbx2* expression counts in the selected Midbrain and Hindbrain regions.

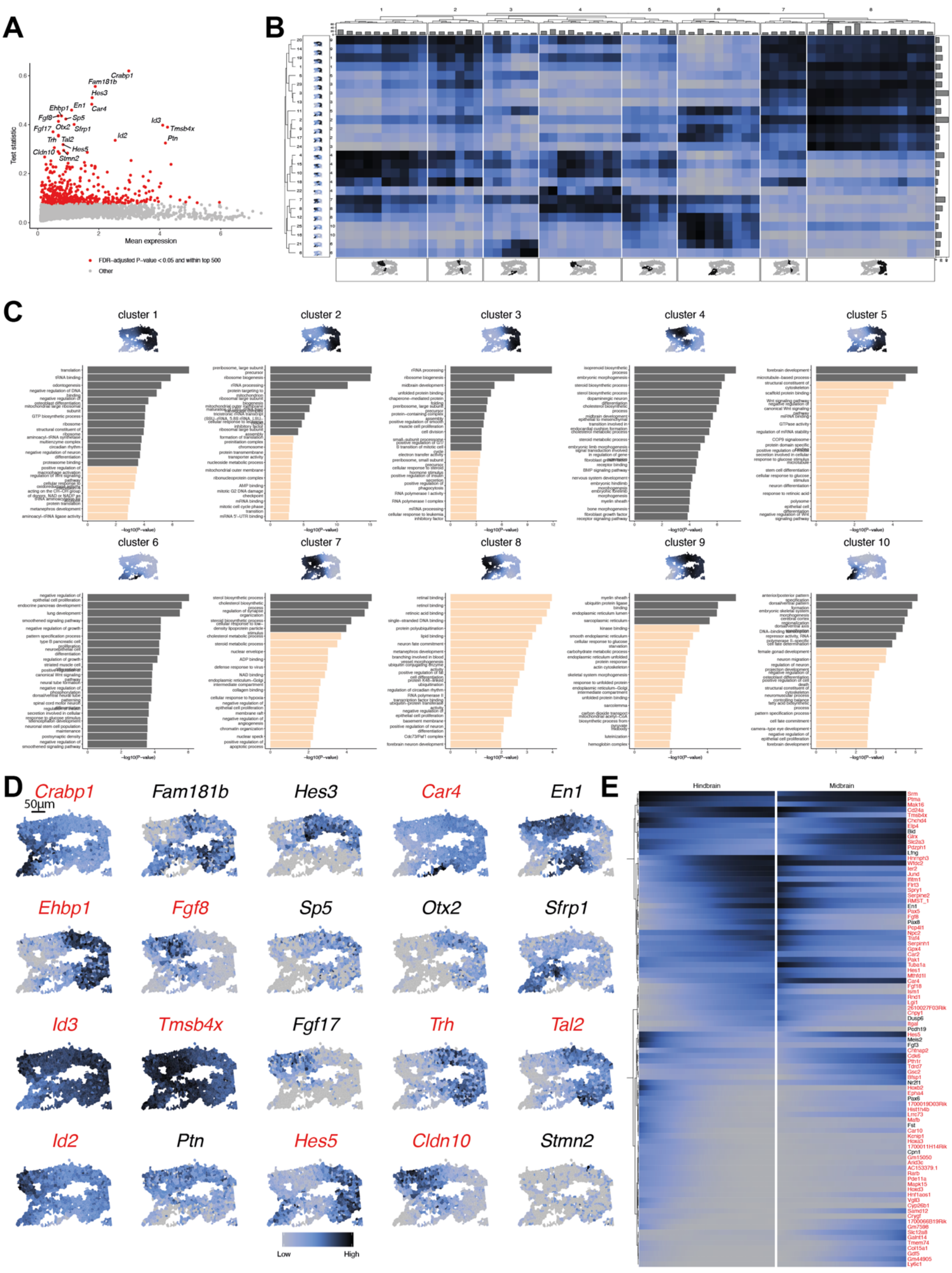


Supp. Figure 20

**Supplementary Figure 20: Statistical interrogation of the Midbrain-Hindbrain Region.**
(A) Scatterplot of all imputed genes, showing mean expression (x-axis) and scHOT weighted mean test statistic (y-axis). Significant (FDR-adjusted P-value $< 0.05$) and top 500-ranked genes are colored red, and the top 20 genes are labeled.

(B) Heatmap of expression of clustered MHB genes and cells, split along columns by clustered cell regions, and along rows by mean expression profiles. Top barplots display the number of cells within each group, right barplots display the number of genes withing each group, bottom spatial graphs display cells belonging to each split cluster, and left spatial graphs show the mean spatial expression for genes that characterize each split cluster.

(C) Spatial graph of the MHB with cells colored by mean expression of the genes belonging to each cluster, and barplots displaying the top 20 enriched gene ontology terms with bar length corresponding to -log10(unadjusted P-value), dark grey bars correspond to FDR-adjusted P-value $< 0.05$.

(D) Spatial graphs of the MHB for the top 20 ranked scHOT weighted mean genes, with red titles corresponding to inferred gene expression.

(E) Smoothed heatmap of cells (columns), ordered along DPT split by anatomical midbrain and hindbrain regions, for genes strongly correlated with DPT (rows). Cells are ordered from low to high DPT from left to right for the hindbrain region, and ordered from high to low DPT from left to right for the midbrain region. Gene names in red correspond to inferred gene expression.

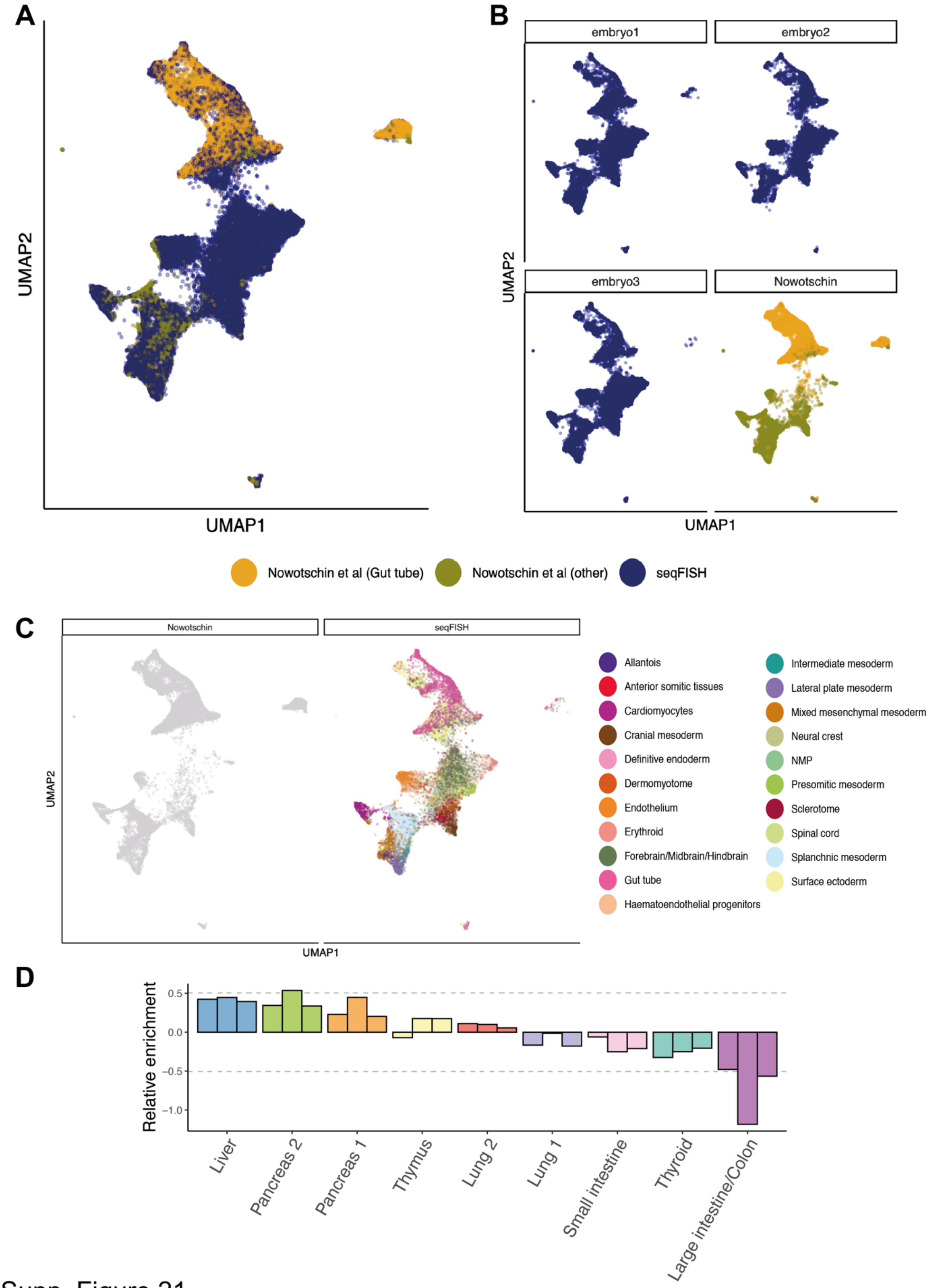


Supp. Figure 21

**Supplementary Figure 21: Integration with Nowotschin *et al.* data.**

(A) Joint UMAP of Nowotschin *et al.* and seqFISH expression data, with cells colored by dataset, and for the Nowotschin *et al.* dataset, whether the cell has an associated developing gut tube cell annotation.

(B) Joint UMAP of Nowotschin *et al.* and seqFISH expression data, with panels corresponding to each embryo and the Nowotschin *et al.* dataset. Colors as in A.

(C) Joint UMAP of Nowotschin *et al.* and seqFISH expression data, where seqFISH cells are colored by their refined cell type annotations based on integration with the Gastrulation atlas dataset.

(D) Barplot of relative enrichment in abundance of seqFISH cells compared to Nowotschin *et al.* cells, each bar corresponds to embryo 1, 2, and 3, from left to right.

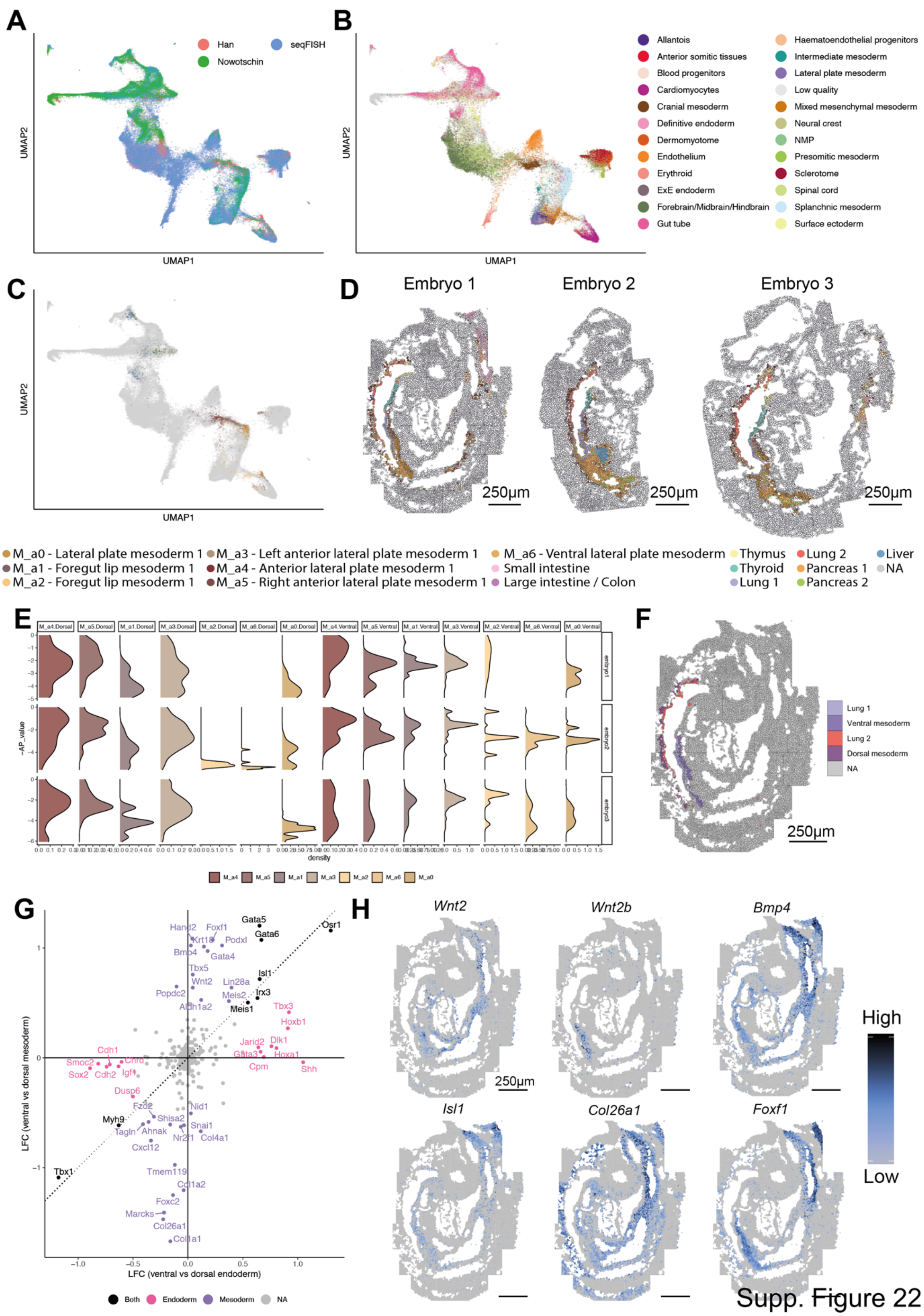


Supp. Figure 22

**Supplementary Figure 22: Surrounding mesoderm of the developing gut tube**

(A) Joint UMAP of Nowotschin *et al.*, Han *et al.* and seqFISH expression data, with cells colored by dataset.

(B) as in (A) with cells colored by corresponding Gastrulation atlas cell type (automatically inferred for cells not coming from the seqFISH dataset).

(C) as in (A) with cells colored by mesodermal and endodermal subtype for the Han *et al.* dataset, and all other cells colored in grey.

(D) Spatial graphs of gut tube and surrounding mesodermal cells, colored by inferred gut tube subtype and mesodermal subtypes respectively.

(E) Density graphs of seqFISH mesodermal cells ordered along physical anterior to posterior axis, split by embryo (rows), and mesoderm cluster and position along dorsal-ventral axis (columns).

(F) Spatial graph of cells corresponding to gut tube subtypes Lung 1 and Lung 2, as well as surrounding mesodermal cells.

(G) Scatterplot of log-fold changes corresponding to tests for differential expression between ventral (Lung 1) and dorsal (Lung 2) endodermal cells (x-axis), and ventral and dorsal mesodermal cells (y-axis) for all seqFISH genes. Significant (FDR-adjusted P-value $< 0.05$ and absolute LFC $> 0.2$) genes are labeled, and colored according to the comparison in which they are selected.

(H) Spatial graphs of expression of selected genes among those differentially expressed between dorsal and ventral subgroups.

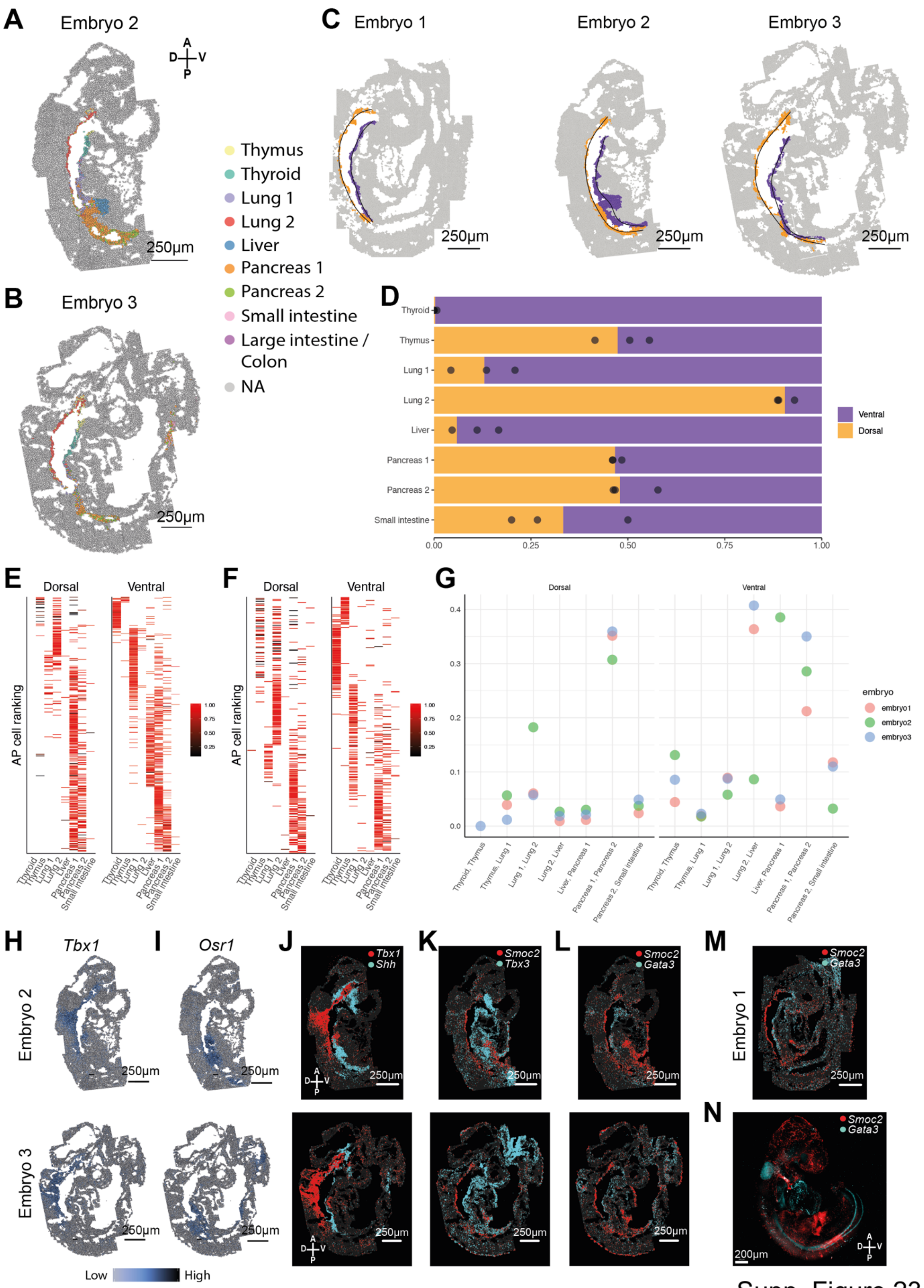


Supp. Figure 23

**Supplementary Figure 23: Comparison between dorsal and ventral side of developing gut tube.**

(A) Spatial map of cells corresponding to the developing gut tube for embryo 2. Scale bar 250 µm.

(B) as in A, for embryo 3.

(C) Spatial map of anatomical foregut cells for embryos 1, 2, and 3, virtually dissected to correspond to the dorsal (orange) and ventral (purple) regions of the developing gut tube. Black lines correspond to the fitted principal curve model for each embryo and developing gut tube region, where cells are ordered from anterior to posterior using these models. Scale bars 250 µm.

(D) Barplot showing relative proportion of cells in ventral or dorsal anatomical region of the developing hindgut, split by classification of developing gut tube subtype. Black points correspond to relative proportions for each individual embryo.

(E) Anterior-posterior ranking of embryo 2 cells, corresponding to each gut tube subtype, split into dorsal and ventral regions. Bar color corresponds to the mapping score associated with classification into the subtype.

(F) as in E for embryo 3.

(G) Scatterplot of anterior-posterior logistic regression prediction error rate (y-axis) for each contiguous pair of developing gut tube subtypes (x-axis), split into dorsal and ventral anatomical regions, for each embryo. A higher prediction error rate corresponds to a higher level of relative mixing of subtypes along the anterior-posterior axis, while a lower prediction error rate corresponds to more distinct and separate arrangement of subtypes along the anterior-posterior axis.

(H) Spatial expression of *Tbx1* only in the developing gut tube for embryos 2 (top) and 3 (bottom). Scale bar 250 µm.

(I) as in H for gene *Osr1*.

(J) '*Digital in situ'* showing detected mRNA molecules for *Tbx1* (red) and *Shh* (cyan) for embryos 2 (top) and 3 (bottom). Scale bar 250 µm.

(K) as in J for genes *Smoc2* (red) and *Tbx3* (cyan).

(L) as in J for genes *Smoc2* (red) and *Gata3* (cyan).

(M) ‘Digital *in situ*’ showing detected mRNA molecules for *Smoc2* (red) and *Gata3* (cyan) for embryo 1. Scale bar 250 µm.

(N) Multiplexed mRNA imaging of whole-mount E8.75 mouse embryo using hybridization chain reaction (HCR) of *Smoc2* (red) and *Gata3* (cyan).

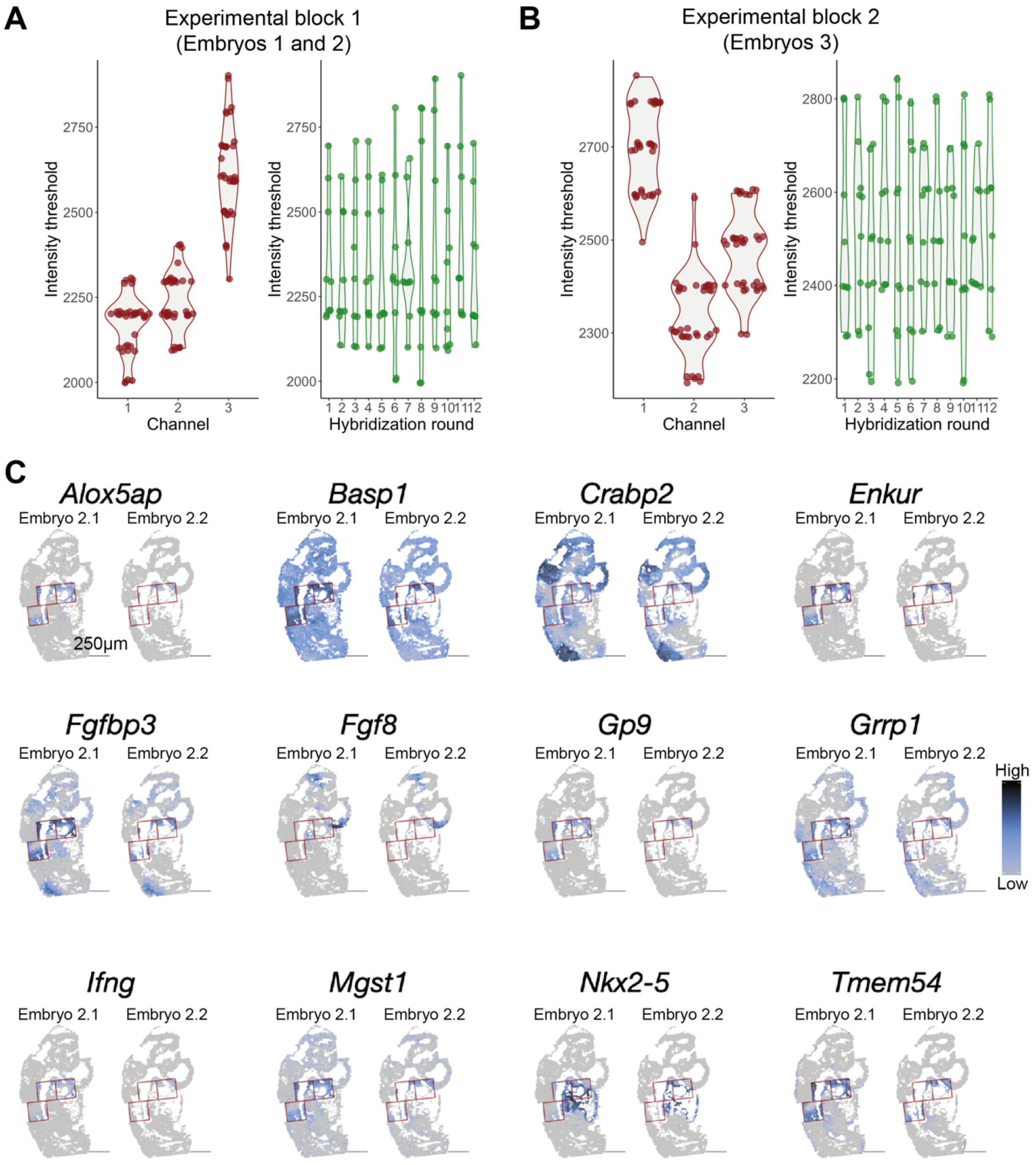


Supp. Figure 24

**Supplementary Figure 24: Channel effect on the distribution of background noise for non-barcoded smFISH data.**

(A) The intensity threshold that separates background spots is highly dependent on the channel (fluorescence) a gene was probed with, but not on the hybridization round. Violin plots on the left show intensity thresholds for experimental block 1 for each gene and field of view combination grouped by color channel, while violin plots on the right show intensity thresholds for each gene and field of view combination grouped by hybridization round.

(B) as in A for experimental block 2 (embryo 3).

(C) Spatial maps of non-barcoded smFISH genes that were probed with Alexa Fluor 647 (AF647) for embryo 2, with red squares around fields of view 39, 40, and 44, which display a strong field of view effect, regardless of the choice of intensity threshold.

### SUPPLEMENTARY TABLES

**Supplementary Table 1: Primary probes**

List of primary probes for seqFISH library, non-barcoded sequential smFISH genes and Eef2 probeset A and B. Each entry contains a primary probe sequence or a primary probe sequence combined with readout probe sequences.

**Supplementary Table 2: Readout probes and decoding strategy for seqFISH probe library**.

List of readout probe sequences and the corresponding fluorophore conjugated to the probes used in the seqFISH experiment. Additionally, a list assigning a unique combination of four pseudocolors and the corresponding readout probe sequence to each gene of the seqFISH library is included; this allows decoding of the barcodes over the multiple imaging rounds.

**Supplementary Table 3: Readout probes for non-barcoded sequential smFISH genes**

List of readout probe sequences and the corresponding fluorophore used for each gene measured by non-barcoded sequential smFISH.

**Supplementary Table 4: List of spatial heterogeneity test results for each of the assigned cell types.**

Columns correspond to gene name, proportion of variability explained by neighboring genes' expression, P-value, t-statistics, and FDR-adjusted P-value.

**Supplementary Table 5: List of spatial heterogeneity test results for each of the Forebrain/Midbrain/Hindbrain subclusters**

Columns correspond to gene name, proportion of variability explained by neighboring genes' expression, P-value, t-statistics, and FDR-adjusted P-value.

**Supplementary Table 6: List of significantly differentially expressed genes between virtually dissected midbrain and hindbrain regions of embryo 2.** Columns correspond to gene, FDR-adjusted P-value, log-fold change for midbrain, mean expression and direction of significance.

**Supplementary Table 7: Top 500 spatially variable genes in the virtually dissected midbrain/hindbrain region of embryo 2**. Columns correspond to gene, mean gene expression across all cells, scHOT weighted mean test statistic, FDR-adjusted P-value, significance ranking, gene cluster cutting hierarchical clustering tree for 25 clusters, and for 10 clusters.

**Supplementary Table 8: List of differentially expressed genes between Lung1 and Lung2 subcluster.** Columns correspond to gene name, P-value, FDR-adjusted P-value, log-fold change for Lung 1 / Lung 2, mean expression of cells in Lung 1 group, and in Lung 2 group.
